# PAPγ associates with PAXT nuclear exosome to control the abundance of PROMPT ncRNAs

**DOI:** 10.1101/2023.10.04.560886

**Authors:** Xavier Contreras, David Depierre, Charbel Akkawi, Marina Srbic, Marion Helsmoortel, Olivier Cuvier, Rosemary Kiernan

## Abstract

Pervasive transcription of the human genome generates an abundance of RNAs that must be processed and degraded. The nuclear RNA exosome is the main RNA degradation machinery in the nucleus. However, nuclear exosome must be recruited to its substrates by targeting complexes, such as NEXT or PAXT. By proteomic analysis, we have identified additional subunits of PAXT, including many orthologs of MTREC found in *S. pombe*. In particular, we show that polyA polymerase gamma (PAPγ) was associated with PAXT. Genome-wide mapping of the binding sites of ZFC3H1, RBM27 and PAPγ, showed that PAXT is recruited to the TSS of hundreds of genes. Loss of ZFC3H1 abolished recruitment of PAXT subunits including PAPγ to TSSs and concomitantly increased the abundance of PROMPTs at the same sites. Moreover, PAPγ, as well as MTR4 and ZFC3H1, was implicated in the polyadenylation of PROMPTs. Our results thus provide key insights into the direct targeting of PROMPT ncRNAs by PAXT at their genomic sites.

## Introduction

Transcription is an essential process that allows production of RNA from the DNA genome. However, cryptic transcription, including antisense transcription, leads to the production of many transcripts that must be degraded^1–3^. Nuclear exosome is the main machinery involved in processing and degrading these cryptic transcripts. One such type of transcript that are prominent and widely studied are promoter upstream transcripts (PROMPTs), also known as upstream antisense RNAs (uaRNAs)^4^.

PROMPTs are short transcripts of about 500-2000 nucleotides in length that initiate upstream and antisense from the TSS of genes. PROMPTs must be eliminated for normal cell function. Indeed, inhibiting the degradation of PROMPTs can lead to deleterious side effects such as the inhibition of translation of mRNAs in the cytoplasm^4^. However, recent data have shown that PROMPTs are required for the transcriptional activation of estrogen-responsive genes through activation of 7SK-P-TEFb complex^5^ suggesting that the abundance of PROMPTs should be tightly regulated. Degradation of PROMPTs is carried out by the nuclear RNA exosome, a highly conserved 3’–5’ ribonucleolytic complex^6–9^. The nuclear exosome consists of 9 core subunits that form a barrel-like structure with a central channel^10,11^. The catalytic activity is conferred by association with 3’-5’ ribonucleases, Rrp6, which possesses distributive activity, and Dis3, a processive exo- and endonuclease^12–17^. Nuclear RNA exosome also associates with MTR4 (SKIV2L2), a 3′-5′ DExH-box RNA helicase that promotes unwinding and degradation of structured RNA substrates^18–20^.

Nuclear exosome utilizes adaptor complexes in order to target RNAs for degradation. The nuclear exosome targeting (NEXT) complex consists of MTR4 together with the zinc-finger containing protein, ZCCHC8, and an RNA-binding protein, RBM7. NEXT targets primarily short, mono-exonic RNAs, such as enhancer RNAs (eRNAs) and PROMPTs/uaRNAs, as well as long non-coding RNAs (lncRNAs), without a requirement for polyadenylation^21,22^. A second exosome targeting complex, consisting of MTR4 together with the zinc-finger protein ZFC3H1, is known as the polysome protector complex (PPC) or the polyA-tail exosome targeting (PAXT) connection^4,23^. PPC/PAXT consists of MTR4, ZFC3H1 zinc-finger protein, ZC3H3, RBM26/27 together with polyA binding protein, PABPN1^23–25^. The presence of PABPN1 confers a preference for polyadenylated RNAs as targets of PAXT. Indeed, PAXT and NEXT share many target RNAs, including PROMPTs/uaRNAs, eRNAs and prematurely terminated sense transcripts, although targeting likely occurs at different stages of maturation^4,23^. PAXT subunits co-localize with pA(+) RNA foci, whose formation depends on ZFC3H1^24^.

While the addition of a poly(A) tail is essential for normal mRNA biogenesis, polyadenylation can also stimulate degradation of aberrant mRNAs and certain ncRNAs, including PROMPTs^26–30^. The canonical mammalian poly(A) polymerases, PAPα and PAPγ, catalyze template-independent polyA extension of the 3’ end of RNA^31–33^. PAPα and PAPγ have similar organization of structural and functional domains^31–34^. PAPγ shares greater than 60% identity to the well-characterized PAPα at the amino acid level and is thought to have arisen by gene duplication of the latter^31,32^. However, PAPγ is localized exclusively in the nucleus while PAPα exhibits both nuclear and cytoplasmic localization. The difference in subcellular localization is thought to be due to signals present in the unique C-terminal region (amino acids 507-736) of PAPγ^31–33^. PAPγ carries out both nonspecific and CPSF/AAUAAA-dependent polyadenylation activity. The catalytic efficiency of PAPγ is similar to that of PAPα. PABPN1 acts as a coactivator of both PAPα and PAPγ. Indeed, PABPN1 plays a role in RNA polyadenylation by strongly increasing the processivity of poly(A) polymerases, leading to hyperadenylation of RNA targets with the addition of up to 800 adenosines^26,27,29^. Interestingly, the nuclear exosome has been shown to be involved in PABPN1 and PAP-mediated decay of intronless β-globin and PANΔENE reporters^26^.

TRAMP is a nuclear polyadenylation complex that was initially characterized in *Saccharomyces cerevisiae*^19,35^. It is composed of a non-canonical PAP, TRF4p, together with MTR4p helicase and a zinc knuckle protein, Air2p. TRF4p adds a short polyA tag to the 3’ end of the target RNA, which is required for its degradation by the nuclear exosome^19^. Such a mechanism was suggested for the NEXT complex via PAPD5/ZCCHC7 subunits^22^. However, it is unclear how PAXT substrates become polyadenylated. PAXT substrates harbor a long polyA tail that is bound by PABPN1 to facilitate recruitment of PAXT. In addition, PROMPTs were resistant to degradation by the exosome when cells were treated using cordycepin, an inhibitor of polyadenylation, highlighting the importance of the polyA tail for degradation. However, the specific polyA-polymerase involved has not been described.

*S. Pombe* Mtl1-Red1 Core (MTREC) is an 11-subunit complex that is thought to be homologous to PAXT^36^. MTREC consists of several modules that are bridged by Red1, which acts as a scaffold. Red1 has been proposed to be the homolog of human ZFC3H1. MTR4, and ZC3H3 are the homologs of MTL1 and Red5, respectively, while both RBM27 and RBM26 are homologs of Rmn1. Interestingly, MTREC contains a polyA binding protein, Pab2 that is a homolog of PABPN1, and a polyA polymerase, Pla1, for which the canonical polyA-polymerases, PAPα and PAPγ, have been proposed as homologs^37^. To date, no direct link between a polyA polymerase and PAXT complex has been described. However, it was shown that both PABPN1 and canonical polyA-polymerases are involved in the decay of mRNA, such as those with retained introns, and PROMPTs. However, no distinction was made between the canonical polyA polymerases, PAPα and PAPγ^26,27^.

In this study, we sought to address the mechanisms underlying the processing of RNA targets by PAXT. Using proteomics of immunopurified ZFC3H1, we identified the polyA polymerase PAPγ as the sole PAP detected in the PAXT complex. We mapped for the first time the localization of subunits of PAXT on chromatin using ChIP-seq. We found that PAPγ co-localizes with PAXT subunits, ZFC3H1 and RBM26, at the TSS of hundreds of genes. Importantly, ZFC3H1 is required for PAPγ recruitment at these sites. RNA-seq analysis showed that loss of ZFC3H1 or PAPγ was associated with the accumulation of PROMPTs at genes directly targeted by PAXT. We further showed that PAXT, including PAPγ, is implicated in the processing and polyadenylation of PROMPTs. Thus, we demonstrate that PAPγ associates with PAXT and is essential for polyadenylation and subsequent degradation of PROMPTs. This study uncovers a connection between the nuclear polyA polymerase, PAPγ, and the PAXT complex that contributes to the processing of PROMPTs. It further indicates that PAXT substrates such as PROMPTs are likely targeted and possibly degraded at their site of production on chromatin.

## Results

### PAPγ is a component of the PAXT complex

In order to gain insight into PAXT function, we first created a HEK-293T cell line expressing ZFC3H1 with an N-terminal Flag-HA (FH) tag using Crispr-Cas9 technology to insert the tag at the endogenous locus. FH-ZFC3H1 interacted with a known ZFC3H1 partner, MTR4, as expected (Supplementary Fig.1A). FH-ZFC3H1 was purified from nuclear extracts by tandem affinity purification. Compared to control cells, a band at the expected size for FH-ZFC3H1 could be distinguished in a silver-stained gel (Supplementary Fig.1B). Analysis of tandem affinity purified FH-ZFC3H1 by mass spectrometry identified 131 interactants having 2 or more peptides and FC>2 compared to the control HEK-293T sample. Among the most represented pathways obtained using gene ontology analysis were RNA processing and splicing (Supplementary Fig. 1C). Polyadenylation was also significantly overrepresented, which might be expected given the connection of PAXT with polyadenylation. Among the interactants, ZFC3H1 and exosome subunits such as MTR4, but not ZCCHC8, were detected (Supplementary Table S1), which is consistent with a previous study^23^. Interestingly, all proposed human orthologs of the yeast MTREC complex were strongly represented among ZFC3H1 interactants (Fig.1a,b). Red1 found in S. pombe has been proposed to be the homolog of human ZFC3H1. MTR4, ZC3H3 and PABPN1 are the homologs of Mtl1, Red5 and Pab2, respectively, while both RBM26 and RBM27 are homologs of Rmn1. Interestingly however, the sole homolog identified for the polyA polymerase Pla1 was PAPγ, a strictly nuclear canonical polyA polymerase. Notably, although PAPα and PAPγ share redundant functions^26,27^, the ZFC3H1 interactome contained only PAPγ (Supplementary Table S1). PAPα was not detected among ZFC3H1 interactants.

In order to confirm interactions, we next performed co-immunoprecipitation analysis of endogenous, unmodified proteins present in HeLa nuclear extracts (Fig. 1c and Supplementary Fig. 2A). As expected, ZFC3H1 was found to interact with subunits of PAXT, such as MTR4, RBM26 and RBM27, as well as PABPN1. Notably, the interaction between ZFC3H1 and PAPγ was confirmed by co-immunoprecipitation. PAPγ was furthermore found to interact with PAXT subunits RBM26, RBM27 and PABPN1 (Supplementary Fig. 2A). RBM26 and RBM27 were also shown to interact with PAXT subunits (Supplementary Fig. 2A). In addition to PAPγ and PABPN1, proteomic analysis identified other ZFC3H1 interactants involved in polyA processing, including CPSF6 and a polyadenosine RNA-binding protein, ZC3H14, also known as MSUT2. Their interaction with ZFC3H1 was confirmed by co-immunoprecipitation (Fig. 1c and Supplementary Fig. 2A). Although Mmi1 and Iss10 homologs, YTHDC1 and YTHDC2, were not identified among the interactants, a previous study identified YTHDC1 as an interactant of ZFC3H1 upon over-expression^23^. Co-IP analysis confirmed that both m6A readers, YTHDC1 and YTHDC2, interacted with endogenous PAXT subunits, including PAPγ (Fig. 1c and Supplementary Fig. 2A). To determine whether PAXT subunits interact in an RNA-dependent manner, we performed co-immunoprecitation analysis in the presence and absence of RNAse. Notably, interaction of PAPγ with ZFC3H1, MTR4 and RBM27 was independent of RNA, whereas interaction between PAPγ and PABPN1 was found to be RNA-dependent. Similar results were obtained for ZFC3H1 (Supplementary Fig. 2B). Taken together, these data identify additional partners of the ZFC3H1 subunit of the PAXT complex. In particular, the proposed orthologs of yeast MTREC subunits were identified and/or confirmed by co-immunoprecipitation as interactants of endogenous ZFC3H1.

**Fig. 1.**
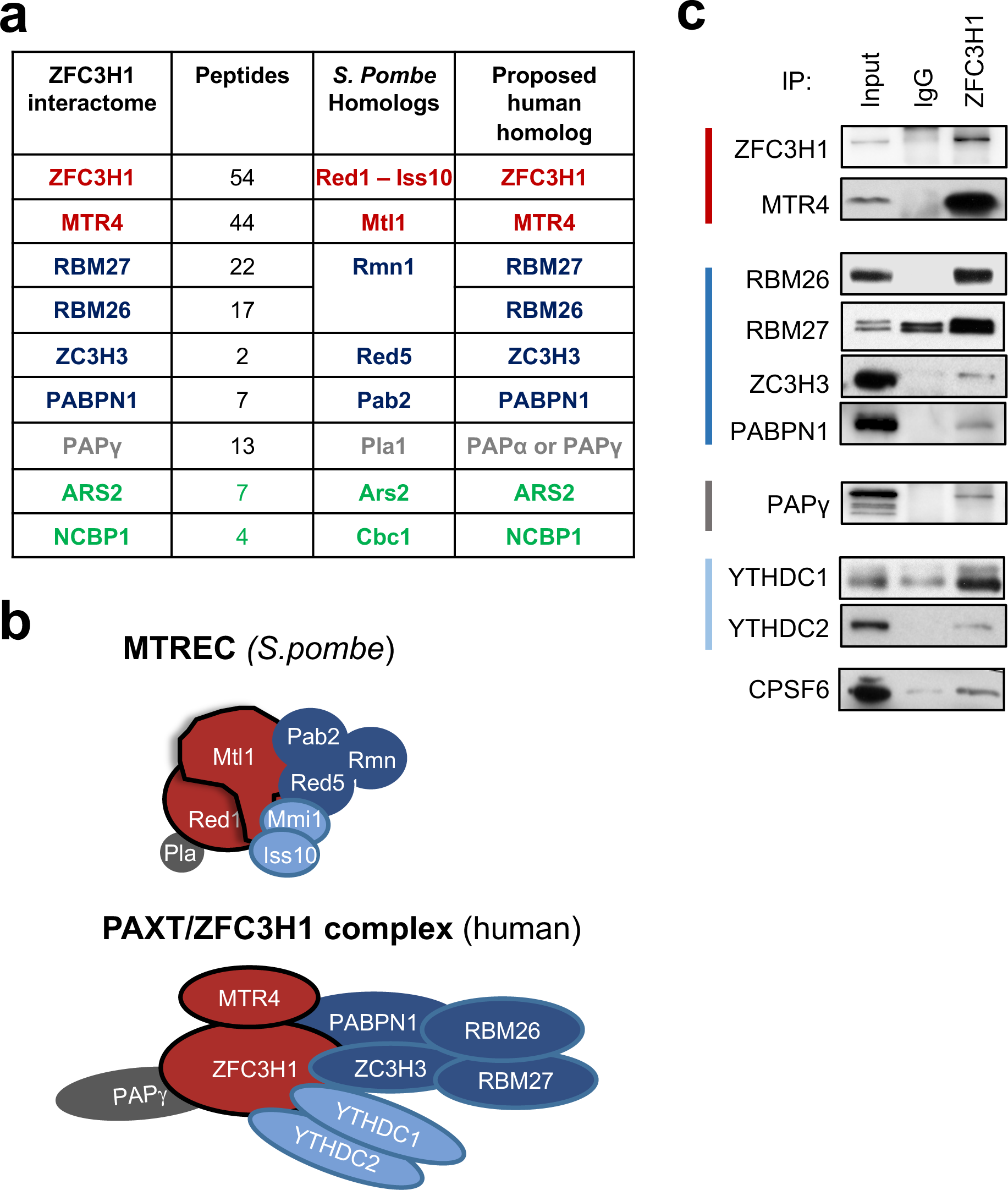
PAPγ is a component of the PAXT complex. **a** Table showing MTREC subunits found in *S. pombe*, the proposed human homologs, the corresponding proteins identified in the ZFC3H1 interactome together the number of unique peptides identified by mass spectrometry. **b** Schematic model depicting the MTREC complex in *S. pombe* (top) and the human PAXT complex based on proteins identified in the ZFC3H1 interactome. **c** Co-immunoprecipitation analysis of ZFC3H1. HeLa nuclear extracts were immunoprecipitated using antibodies against ZFC3H1 or an IgG control. Immunoprecipitates and an aliquot of nuclear extract (input, 5%) were analysed by SDS-PAGE followed by immunoblot using the antibodies indicated on the figure.

### PAXT subunits are co-recruited to chromatin

RNA processing, including polyadenylation, frequently occurs co-transcriptionally^38^ and the recruitment of RNA binding proteins, such as splicing factors, can be monitored by chromatin immunoprecipitation (ChIP)^39^. Consistent with this, cellular fractionation analysis revealed that PAXT subunits, MTR4, ZFC3H1, RBM26, RBM27 and PAPγ, partially localized to chromatin in addition to the nucleoplasm (Supplementary Fig. 3A). To better characterize PAXT and its functions, we sought to identify sites of recruitment of some of its key subunits on chromatin. ChIP-seq was performed using antibodies recognizing known PAXT subunits, ZFC3H1 and RBM26, as well as PAPγ, in addition to RNA polymerase II (RNAPII). It should be noted that antibodies against RBM27 did not yield analyzable signal. ChIP-seq reads of ZFC3H1, RBM26 and PAPγ were detected mostly over gene bodies and transcription start sites (TSS) as well as at enhancers. The distribution was similar to that observed for RNAPII (Supplementary Fig. 3B). At genes, the signal was more intense at TSSs and was characterized by a sharp peak (Fig. 2a and Supplementary Fig. 3C). The specificity of PAPγ ChIP-seq signal was confirmed by RNAi-ChIP-qPCR at the TSS of target genes using the same antibody used in ChIP-seq as well as a second independent antibody recognizing PAPγ (Supplementary Fig. 3D).

**Fig. 2.**
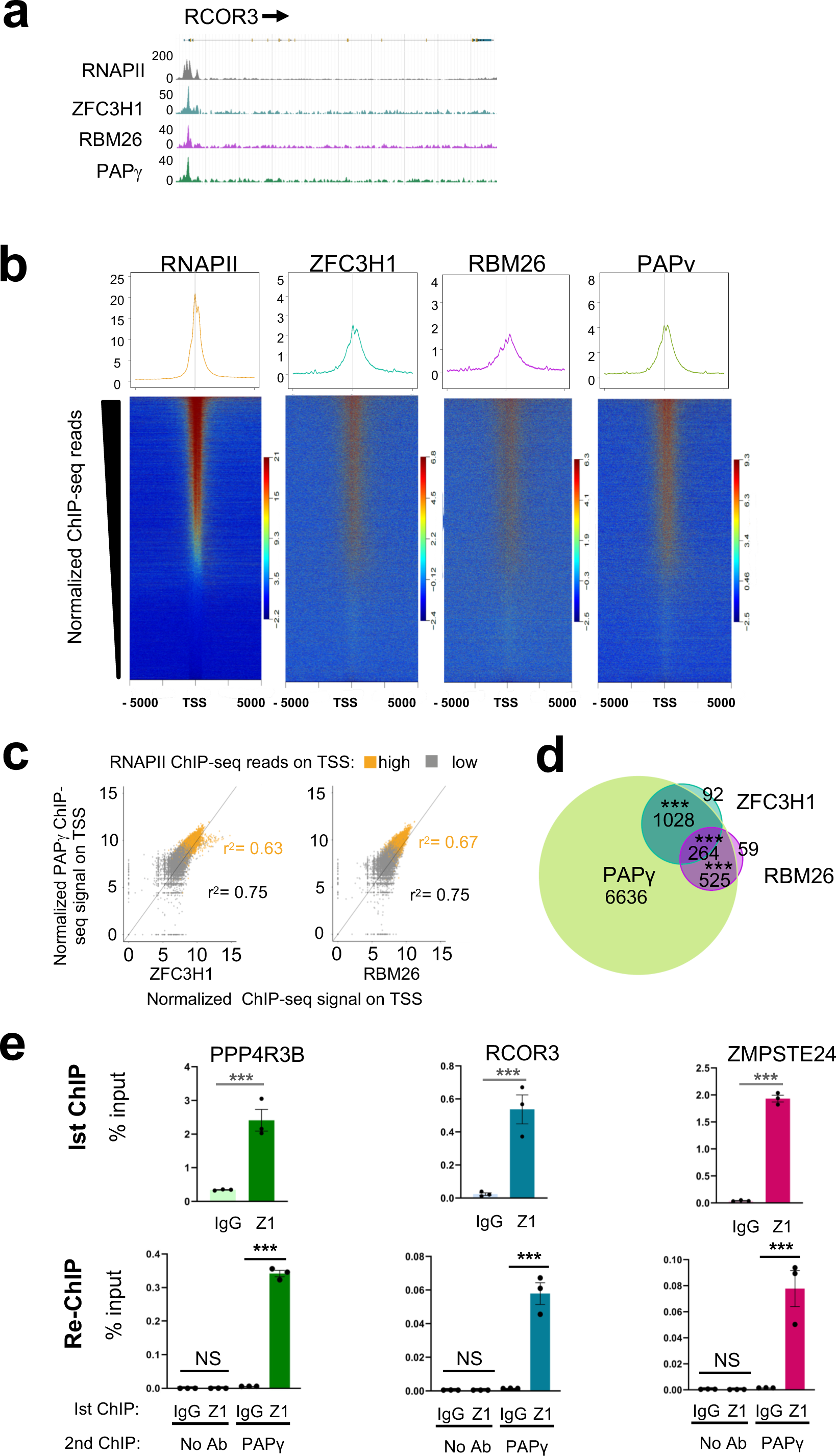
PAXT subunits are co-recruited to chromatin. **a** Browser shots of RNAPII, ZFC3H1, RBM26, and PAPγ ChIP-seq signal over a representative gene in HeLa cells. A schematic representation of the gene is shown above. **b** ChIP-seq heatmaps centered on TSSs ± 5 kb and rank-ordered by normalized RNAPII ChIP-seq signal. Normalized ChIP-seq reads of ZFC3H1, RBM26 or PAPγ were plotted respecting the same ranking. Scaled average density profiles of normalized ChIP-seq reads are shown above. **c** Scatter plots showing the normalized ChIP-seq signal of PAPγ relative to that of ZFC3H1 or RBM26 at the TSS of genes with a high or low occupancy of RNAPII at the TSS, as indicated. **d** Venn diagram showing the overlap of ChIP-seq peaks of ZFC3H1, RBM26 and PAPγ. P-values were calculated using Fisher’s exact test (\*\*\**P* < 0.001). **e** Re-ChIP analysis performed in HeLa cells showing co-localization of ZFC3H1 and PAPγ at the TSS region of the indicated genes. Eluates from an initial ChIP using ZFC3H1, or a control IgG antibody (1st ChIP) were used for PAPγ ChIP or no antibody as a control (re-ChIP). Results of qPCR after 1^st^ and 2^nd^ ChIPs are shown as % of input for 1^st^ ChIP. Data represent mean ± SEM obtained from 3 independent experiments (\*\*\**P* < 0.001, NS indicates not significant, independent Student’s *t* test).

We next addressed whether PAXT subunits bind chromatin as a protein complex. The presence of both ZFC3H1 and PAPγ could be detected at TSSs that are associated with PROMPTs, as shown by ChIP-qPCR (Supplementary Fig. 4A). ZFC3H1, RBM26 and PAPγ ChIP-seq signals were then mapped at TSSs that had been ranked according to RNAPII occupancy. The heatmaps showed that all 3 subunits were associated with an overlapping subset of TSSs that could be largely ranked by RNAPII signal intensity (Fig. 2b). Indeed, binding sites of PAPγ were highly correlated with those of ZFC3H1 or RBM26 as well as with RNAPII (Fig. 2c). Regions bound by ZFC3H1, RBM26 or PAPγ were also bound by RNAPII (Supplementary Fig. 4B) although the correlation with RNAPII occupancy was better for RBM26 and PAPγ compared to ZFC3H1, which may be due to detection thresholds of the different antibodies used.

Further analysis confirmed that peaks of PAPγ, RBM26 and ZFC3H1 also significantly overlapped (Fig. 2d). Indeed, almost all peaks of ZFC3H1 (93,3%, p-value< 1e-79) and RBM26 (93%, p-value< 1e-51) overlapped with peaks of PAPγ. It should be noted however that many peaks of PAPγ did not overlap with those of ZFC3H1 or RBM26, which could be due to lower detection efficiency of these latter factors due to lower efficiencies of the antibodies in immunoprecipitating such complexes, or the association of PAPγ with complexes independent of PAXT.

To further confirm co-recruitment of ZFC3H1 and PAPγ at TSSs, we performed re-ChIP experiments. TSSs associated with ZFC3H1 (1st ChIP) were found to be associated with PAPγ (re-ChIP), demonstrating that both ZFC3H1 and PAPγ are co-recruited at these sites (Fig. 2e). Thus, these results show that PAXT subunits ZFC3H1, RBM26 and PAPγ are recruited to chromatin, at TSSs as well as at enhancers, whose associated RNAs have been shown to be exosome substrates^40,41^. These data furthermore reveal that ZFC3H1, RBM26 and PAPγ frequently co-localize on chromatin, likely as the PAXT complex, at sites of active transcription.

To further investigate recruitment of PAXT to chromatin, we depleted a major targeting subunit, ZFC3H1, and performed ChIP-seq experiments for PAXT subunits. Immunoblot analysis of extracts confirmed depletion of ZFC3H1 but no significant effect on expression of either RBM26 or PAPγ was detected (Supplementary Fig. 5). ZFC3H1 depletion was accompanied by a significant decrease of ZFC3H1 at chromatin, particularly at TSSs (Fig. 3a, top panel, Fig. 3b, left panel, Fig 3c, left panel p-value< 1e-16, Wilcoxon test). Heatmaps of ChIP-seq signal loss over TSSs showed that loss of RBM26 and PAPγ was largely correlated to loss of ZFC3H1. Notably, loss of ZFC3H1 was accompanied by a significant decrease in signal of both RBM26 and PAPγ at TSSs (Fig. 3a and Fig. 3b, middle and right panels, Fig. 3c, middle and right panels, p-values < 1e-16, and < 1e-16 respectively, Wilcoxon test). While recruitment of RBM26 and PAPγ at PAXT binding sites appears to be dependent on the presence of ZFC3H1, we cannot exclude that recruitment of these factors at other sites via additional factors, or through indirect mechanisms, may also occur. Taken together, these data suggest that ZFC3H1 is implicated in the recruitment of other components of the PAXT complex at the genome-wide level.

**Fig. 3.**
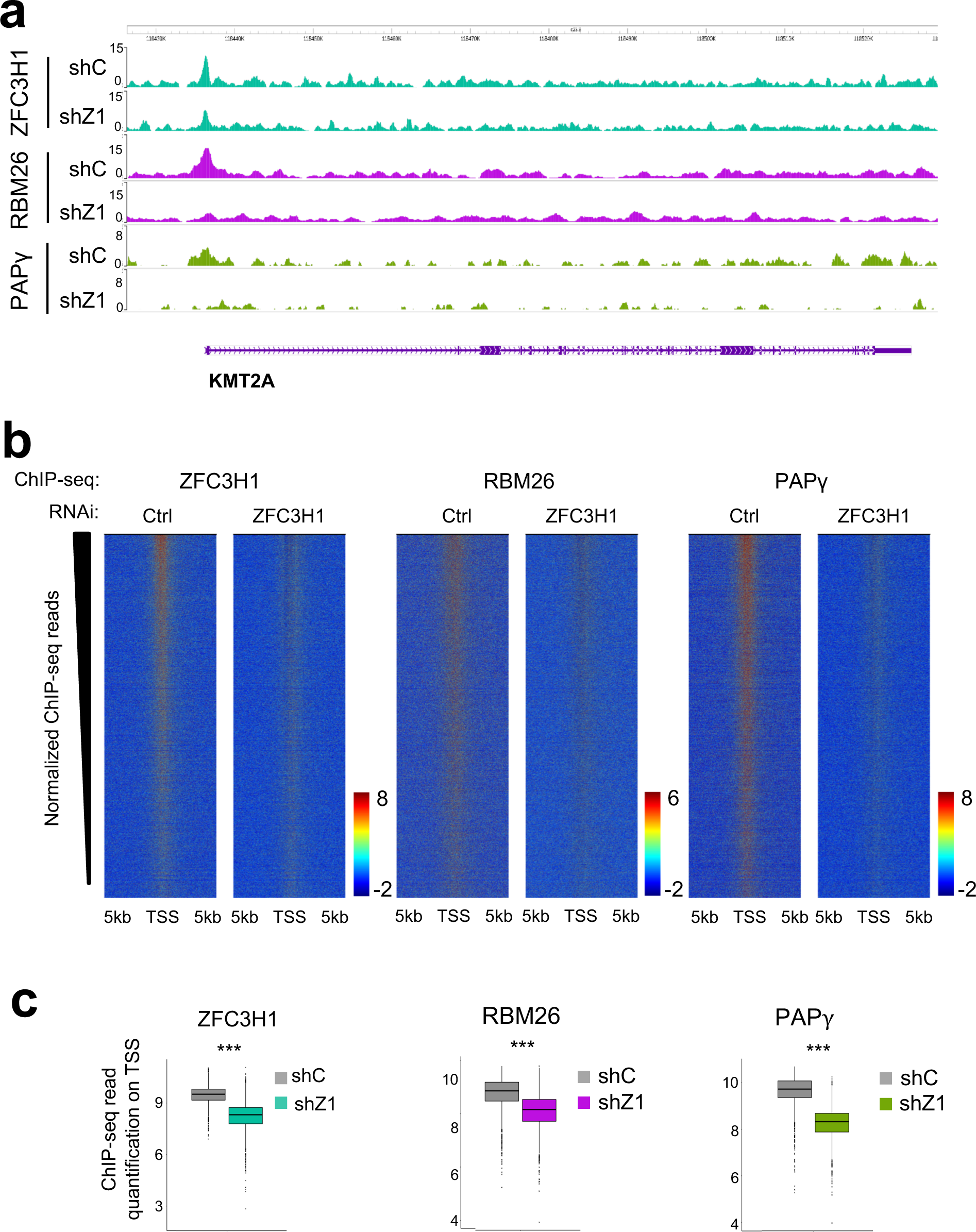
ZFC3H1 is required for recruitment of PAXT to chromatin. **a** Browser shots of ZFC3H1, RBM26, and PAPγ ChIP-seq signal over a representative gene in HeLa cells following loss of ZFC3H1 or a control. A schematic representation of the gene is shown below. **b** Heatmaps centered on TSSs ± 5 kb showing normalized ChIP-seq reads of ZFC3H1, RBM26 or PAPγ in shCon and shZFC3H1 samples and rank-ordered by the change in ZFC3H1 signal between shCon and shZFC3H1 samples. Normalized ChIP-seq reads of RBM26 or PAPγ were plotted respecting the same ranking. **c** Box plot quantification of normalized ChIP-seq reads of ZFC3H1, RBM26 or PAPγ in shCon and shZFC3H1 samples, as indicated, at the top decile of the heat maps shown in b (\*\*\**P* < 0.001, Wilcoxon test).

### PAXT modulates the abundance of PROMPTs

The association of PAXT at TSSs might indicate that the processing of substrates such as PROMPTs occurs in the vicinity of chromatin. To determine if chromatin-association of PAXT is implicated in the processing of PROMPTs genome-wide, we performed *de novo* analysis of RNA-seq data in ZFC3H1 depletion condition^25^. TSSs were oriented 5’-3’ and ranked by the loss of ChIP-seq signal of ZFC3H1, RBM26 or PAPγ, following depletion of ZFC3H1. Change in RNA-seq reads were calculated and mapped at the same sites. Strikingly, the highest accumulation of PROMPTs occurred at TSSs showing the greatest loss of PAXT upon depletion of ZFC3H1 (Fig. 4a,b). To determine if these findings extended to PAPγ, RNA-seq was performed in PAPγ-depleted HeLa cells (Supplementary Fig. 6A) and a similar analysis was carried out. As observed for ZFC3H1, loss of PAPγ also led to the accumulation of TSS-associated RNAs, which could be ranked by the loss of ZFC3H1, RBM26 or PAPγ at the sites following depletion of ZFC3H1 (Fig. 4c). Interestingly, loss of PAPγ did not have a major impact on the abundance of mRNAs (Supplementary Fig. 6B). Only 279 mRNAs were down-regulated and 166 were up-regulated following loss of PAPγ. This finding is consistent with previous studies showing that loss of PAPγ alone did not have a significant effect on RNA abundance ^26,27^. Taken together, these results strengthen the importance of PAPγ and other PAXT subunits in the regulation of PROMPTs. They furthermore show that nuclear exosome-associated factors can be detected at their site of activity, suggesting that targeting and possibly degradation occurs at the site of transcription.

**Fig. 4.**
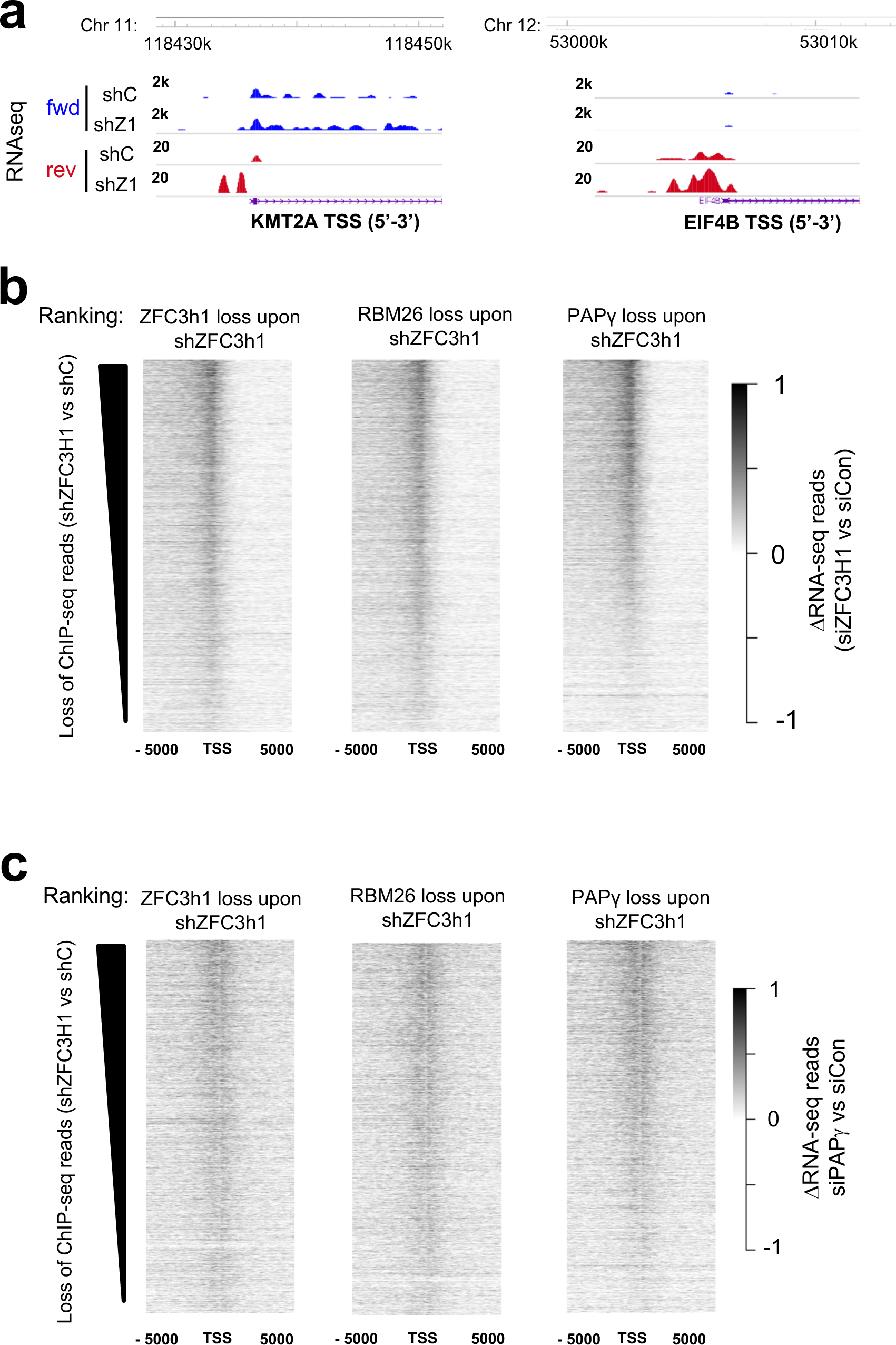
Loss of PAXT subunits leads to the accumulation of PROMPTs genome-wide. **a** Browser shots of RNA-seq reads at the TSS region of representative genes following loss of ZFC3H1 or a control in HeLa cells. A schematic representation of the gene is shown below. **b,c** Heatmaps representing the differential of RNA-seq reads following loss of ZFC3H1 (b) or PAPγ (c) compared to a control knock-down, centered on TSSs ± 5 kb and rank-ordered by change in normalized ChIP-seq reads of ZFC3H1 (left), RBM26 (middle) or PAPγ (right) in HeLa cells.

To further characterize the function of PAPγ and PAXT in the stabilization of PROMPTs, we performed RT-qPCR for PROMPTs associated with TSSs bound by PAXT. As expected, KD of MTR4 as well as that of ZFC3H1 resulted in significant up-regulation of PROMPTs (Fig. 5a and Supplementary Fig. 7A). Loss of PAPγ also increased PROMPTs levels, similarly to depletion of PAXT subunits MTR4 and ZFC3H1. Similar results were obtained using a second independent siRNA targeting either MTR4, ZFC3H1, PAPγ or a non-targeting control (Supplementary Fig. 7B, C).

**Fig. 5.**
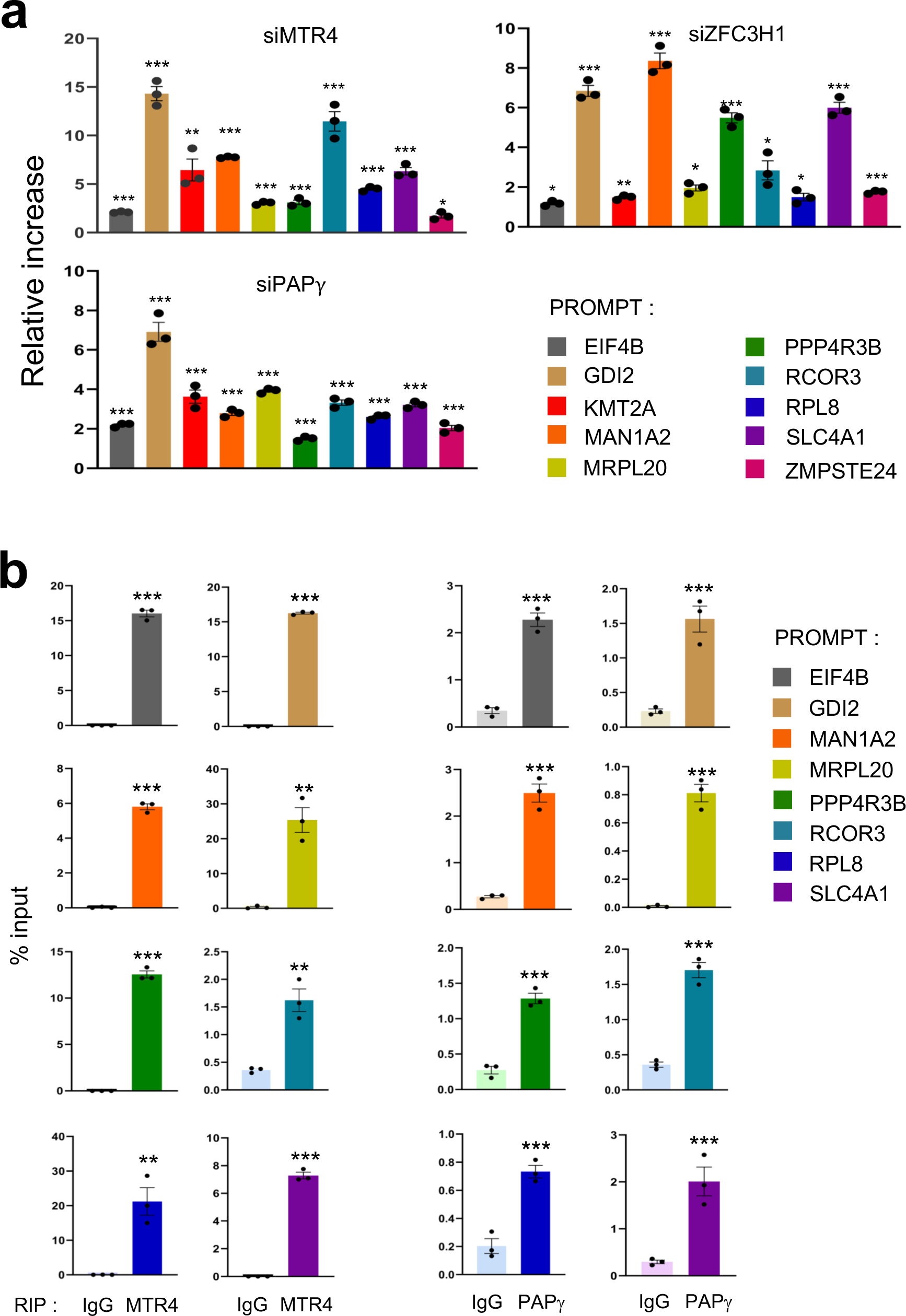
ZFC3H1, MTR4 and PAPγ modulate the abundance of PROMPTs. **a** Total RNA extracts of HeLa cells transfected with siRNAs directed against MTR4, ZFC3H1, PAPγ or a control were analyzed by RT-q-PCR using the oligonucleotide pairs indicated. The values were normalized to those for the control transfection, which was attributed a value of 1. Data represent mean ± SEM obtained from 3 independent experiments (\*\*\**P* < 0.001, \*\**P* < 0.01, \**P* < 0.05, NS indicates not significant, independent Student’s *t* test). **b** Total RNA extracts of HeLa cells were analyzed by RIP using antibody to MTR4, PAPγ or control IgG, as indicated. Immunoprecipitates and an aliquot of extract were analyzed by RT-qPCR using oligonucleotide pairs indicated and the values were expressed relative to the input sample (%). Data represent mean ± SEM obtained from 3 independent experiments (\*\*\**P* < 0.001\*\**P* < 0.01, \**P* < 0.05, independent Student’s *t* test).

To determine whether the PAXT-dependent PROMPTs identified by genome-wide analyses are directly targeted by PAXT complex, we performed RNA immunoprecipitation (RIP) analysis. As shown in Fig. 5b (left panels), PROMPTs were significantly detected in immunoprecipitates of MTR4. Notably, ZFC3H1 co-immunoprecipitated with MTR4, as expected (Supplementary Fig. 7D). Similar results were obtained using anti-PAPγ antibody (Fig 5b, right panels and Supplementary Fig. 7E). These results demonstrate that PAXT subunits, PAPγ and ZFC3H1 together with MTR4, associate with PROMPTs, which likely occurs at the site of transcription, to modulate their abundance. Taken altogether, our data thus support a mechanism for the direct targeting and degradation of PROMPTs by PAXT at sites of ncRNA production on chromatin.

### PAPγ is implicated in the polyadenylation of PROMPTs

Substrates of the PAXT complex are thought to be polyadenylated, as suggested by the association of PABPN1 with PAXT^23^. However, the identity of the polyA polymerase is not known. Since our data show that PAPγ is associated with the PAXT complex and modulates the abundance of PROMPTs, we sought to determine whether PAPγ might polyadenylate PAXT-dependent PROMPTs.

To determine whether PAPγ can polyadenylate PROMPT ncRNAs, we measured polyadenylation by poly(A) test (PAT) assay. The PAT assay detects the polyA tail of a specific RNA target by PCR using an oligodT primer together with a primer containing a sequence near the PAS signal of the target RNA. Polyadenylated RNAs generate PCR products with heterogeneous electrophoretic mobility, appearing as a smear. Extracts of HeLa cells transfected with siRNAs targeting either MTR4, ZFC3H1, PAPγ or a non-targeting control were used in PAT analysis (Fig. 6a). RNAs of several PROMPTs, as well as a control RNA, K-Ras, were first normalized among the different knock-down conditions using PCR primers detecting an internal product (Fig. 6b, bottom panel, F+R). The polyA tail of each PROMPT was then detected using an internal primer together with an oligo (dT) primer (Fig. 6b, top panel, F+T). Knock-down of MTR4, ZFC3H1 or PAPγ significantly reduced the intensity of the polyA tail signal of PROMPTs. In contrast, the intensity of the polyA tail signal of K-Ras mRNA was slightly reduced by depletion of MTR4, and slightly increased by loss of either ZFC3H1 or PAPγ (Fig. 6b, top panel, F+T, Fig. 6c). The polyA tail signal was largely abolished by treatment of the reaction products with RNaseH in the presence of oligo (dT), confirming the specificity of the reaction for polyA polymers (Fig. 6d). Altogether, our data show that PAPγ, which is associated with PAXT complex, contributes to the polyadenylation of PROMPTs.

**Fig. 6.**
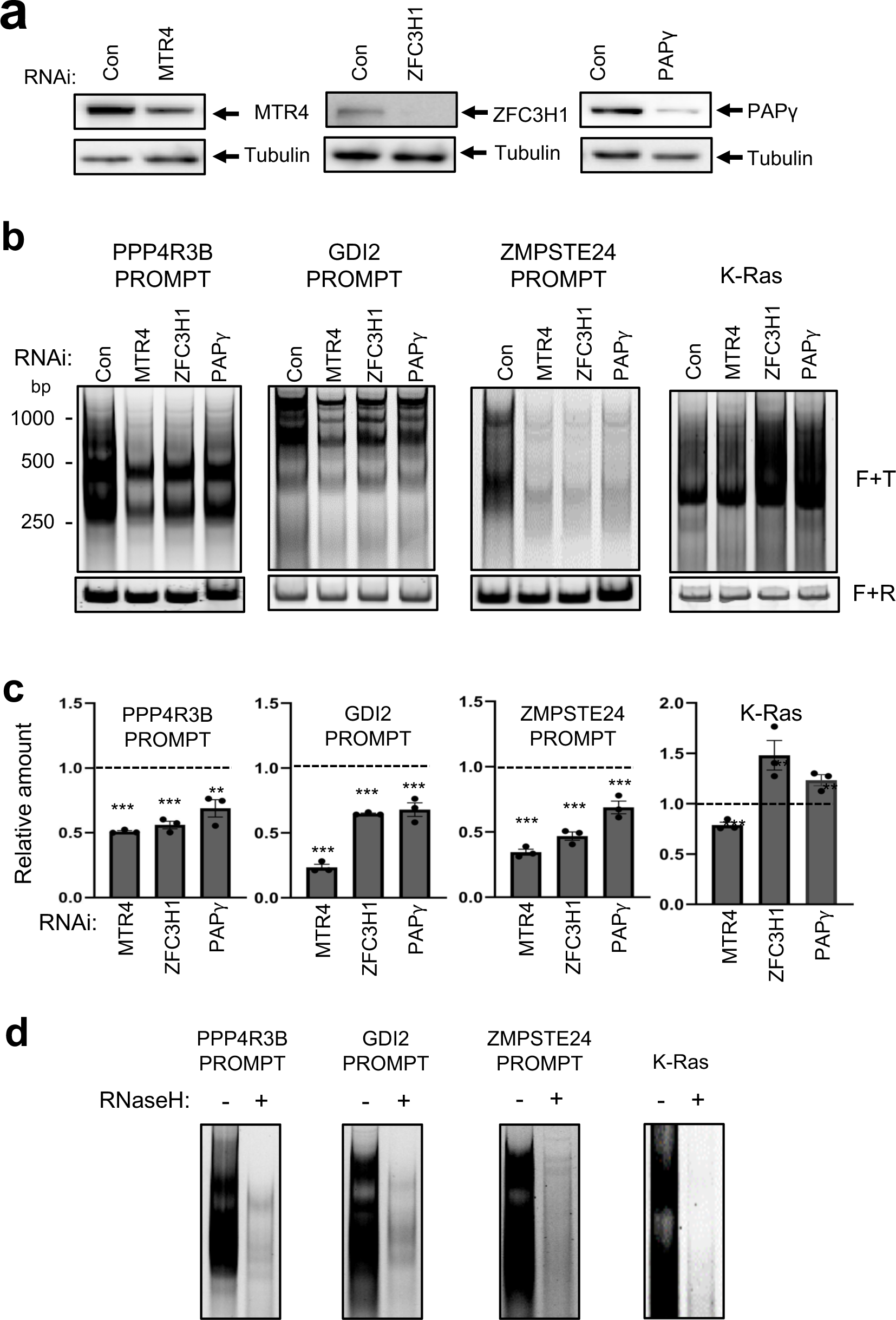
PAPγ is implicated in the polyadenylation of PROMPTs. **a** Immunoblot analysis of HeLa cell extracts following transfection with siRNAs directed against MTR4, ZFC3H1, PAPγ, or a control (con), as indicated on the figure. Samples were analysed by SDS-PAGE followed by immunoblot using the antibodies indicated on the figure. **b** Representative polyacrylamide gels showing electrophoretic mobility of PCR products using extracts of HeLa cells transfected with siRNAs targeting MTR4, ZFC3H1, PAPγ, or a nontargeting control (con). PCRs were performed using a forward primer specific for the gene indicated above the gel together with oligo (dT) (F+T; top panel) or with a specific reverse primer (F+R; bottom panel). Size markers are shown at left. **c** Quantification of F+T signal of gels, as shown in b. Values were expressed relative to the control sample, which was attributed a value of 1. Data represent mean ± SEM obtained from 3 independent experiments (\*\*\**P* < 0.001\*\**P* < 0.01, independent Student’s *t* test). **d** Representative polyacrylamide gels showing electrophoretic mobility of PCR products performed using a specific forward primer together with oligo (dT) (F+T) from extracts of HeLa cells. Samples were treated with RNase H or mock-treated, as indicated on the figure.

## Discussion

The nuclear RNA exosome is the major RNA degradation machinery in the nucleus. However, it must be recruited to its target RNAs by one or more adapter complexes. One such complex, PAXT, recruits nuclear exosome to polyadenylated RNAs, via the PABPN1 subunit. Here, we addressed the mechanisms underlying processing of PROMPTs by PAXT. Using proteomics of purified ZFC3H1, we identified PAPγ, which is a predominantly nuclear polyA polymerase, as the sole PAP detected in the PAXT complex. The localization of subunits of PAXT on chromatin was mapped using ChIP-seq. We found that PAPγ co-localizes with PAXT subunits, ZFC3H1 and RBM26, at the TSS of several hundred genes. Importantly, ZFC3H1 is required for PAPγ recruitment at these sites. Finally, we showed that PAPγ was implicated in the processing of PROMPTs and was required for their efficient polyadenylation. Collectively, these data show that a specific PAP, PAPγ, associates with PAXT and is essential for polyadenylation and subsequent degradation of PROMPTs. These findings uncover a connection between the nuclear polyA-polymerase PAPγ and the PAXT complex that contributes to the processing of PROMPTs. It further indicates that certain PAXT RNA substrates, such as PROMPTs, are targeted and likely degraded at their site of production on chromatin.

Polyadenylation is a major RNA processing step occurring on nascent transcripts that determines the fate of cellular mRNAs (see^42^ for review). In mammals, most mRNAs are polyadenylated, which consists of the addition of ∼250 non-coded adenosines. Polyadenylation confers stability to the mRNA and is required for efficient translation. On the contrary, in prokaryotes, the addition of a polyA tail marks mRNAs for degradation^43–46^. The addition of a short polyA tail (15–40 adenosines) on bacterial transcripts provides a platform for the 3′-exonuclease polynucleotide phosphorylase (PNPase) to initiate 3′–5′ exonucleolytic degradation^47^. Bacterial polyadenylation therefore primarily regulates turnover and quality control of specific cellular transcripts^48^. Polyadenylation of nuclear transcripts could therefore be considered to serve a similar function in mammals^19^. The polyA tail added by the TRAMP complex provides a landing pad for nuclear exosome to facilitate 3’-5’ RNA degradation or trimming^19,49^. However, the TRAMP complex in humans is localized in the nucleolus^22^ and so it was unclear how nucleoplasmic exosome substrates become polyadenylated. PAPγ has been shown to polyadenylate many snRNAs^26,27^, which are also targeted by PAXT^23,50^. Our findings indicate that PAPγ can also polyadenylate PROMPTs that are targeted by PAXT for degradation by nuclear exosome. This finding supports the recent report showing that Pla1 subunit of MTREC hyperadenylates PROMPTs in S. pombe^51^. Interestingly, we observed that loss of the canonical polyA polymerase, PAPα, also affected the abundance of PROMPTs, and showed a cumulative effect in combination with loss of PAPγ (data not shown). However, PAPα did not interact with PAXT in either the presence or absence of PAPγ, suggesting that its effect on PROMPTs does not occur via PAXT. Since concomitant loss of both PAPs has been shown to affect the stability of several RNA species ^26,27^, it will be interesting to determine whether PAPα and PAPγ might co-operate in the hyperadenylation of PROMPTs through PAXT-dependent and -independent mechanisms.

ZFC3H1 can be found in foci in the nucleus together with polyadenylated RNAs^24^, suggesting that degradation of the substrate RNAs may occur in the nucleoplasm. Using ChIP-seq, we found that several subunits of PAXT, such as ZFC3H1, RBM27 and PAPγ, were associated with chromatin at sites of transcription of PROMPTs. This suggests that PROMPTs are likely targeted by PAXT directly at the site of transcription. Whether degradation also occurs on site, or in foci in the nucleoplasm remains unclear.

Polyadenylation and degradation of unstable nuclear RNAs, such as PROMPTs, is important to prevent their deleterious accumulation. Indeed, Manley and colleagues previously showed that polyadenylated PROMPTs and prematurely terminated transcripts that were stabilized by depletion of either MTR4 or ZFC3H1, but not NEXT subunits, accumulated in the nucleus and were also exported to the cytoplasm where they became associated with polysomes^4^. MTR4- or ZFC3H1-depleted cells displayed significant inhibition of translation of mRNAs, likely due to competition with the exported PROMPTs. Thus, the MTR4 and ZFC3H1-containg complex was therefore termed ‘polysome protector complex’ since it assures the proper polyadenylation and subsequent degradation of PROMPTs and prematurely terminated transcripts in the nucleus, thereby preventing their export to the cytoplasm and occupancy of ribosomes^4^. Our data show that loss of PAXT leads to a 2 to 3 fold reduction of polyA tails, as measured by PAT assay, when PROMPTs have been normalized for abundance. However, depletion of PAXT leads to a significant increase in the abundance of PROMPTs in cells, as measured by RT-qPCR. Therefore, it is likely that, although less efficiently polyadenylated, the significant increase in PROMPT abundance is likely sufficient to lead to cytoplasmic export and deregulation of translation as observed previously^4^.

It was previously shown that asymmetric sequence determinants flanking TSSs control promoter directionality by regulating cleavage and polyadenylation of promoter-proximal transcripts^52^. PROMPT/uaRNAs are enriched for PAS signals while being depleted for U1 sites. It was shown that PROMPTs/uaRNAs are cleaved and polyadenylated at poly (A) sites close to the TSS, with a peak of cleavage sites about 700 bp upstream of the TSS, that were associated with the canonical 3’ end processing machinery. Interestingly, a previous study also showed that polyadenylated PROMPTs detected following depletion of MTR4 or ZFC3H1 contain canonical PAS at the 3’ end^4^. Moreover, we confirmed an interaction between ZFC3H1 and CPSF6 subunit of the 3’-end processing machinery. PAPγ is a canonical poly(A) polymerase. It contains a U1 interaction region in its C-terminus and can be inhibited by U1^31^. PAPγ is highly active in both AAUAAA- and CPSF-dependent polyadenylation, as shown using in vitro polyadenylation assays^31,33^. Therefore, the high density of PAS and the relative paucity of U1 sites in PROMPTs would favor polyadenylation of cleaved PROMPTs by PAPγ. Taken together, these findings suggest that the enrichment of PAS in PROMPTs/uaRNAs facilitates transcript cleavage while the association of PAXT with PAPγ subunit of PAXT, which associates with PROMPT RNAs, carries out polyadenylation. PAXT may then target the polyadenylated transcripts to the nuclear exosome for degradation.

Altogether, our data shed light on the processing of PROMPT ncRNAs. PROMPTs were recently shown to be implicated in transcriptional activation at estrogen-responsive genes by regulating transcriptional pause release^5^. Thus, the identification of factors and mechanisms that control the abundance of PROMPT ncRNAs is important to better understand the mechanisms controlling transcription of the coding genome.

## Methods

### Cell culture and reagents

HeLa cells were grown in Dulbecco’s modified Eagle’s minimal essential medium (DMEM) (Sigma-Aldrich, D6429), supplemented with 10% fetal calf serum (FCS; Eurobio Scientific, CVFSVF00-01) and containing 1% penicillin-streptomycin (Sigma-Aldrich, P4333). HEK-293T were grown in Hepes-modified DMEM (Sigma-Aldrich, D6171), supplemented with 10% FCS (Eurobio Scientific, CVFSVF00-01) and containing 1% penicillin-streptomycin (Sigma-Aldrich, P4333). All cells were grown in a humidified incubator at 37°C with 5% CO2.

### Antibodies

Antibodies used in this study are shown in Table S2.

### CRISPR-Cas9 Mediated Editing of endogenous ZFC3H1 gene

An sgRNA targeting the ZFC3H1 gene around the ATG translation start site was cloned in pSpCas9 (BB)-2A-GFP plasmid (Addgene #48138). The plasmid was then transfected into HEK-293T cells along with a single stranded oligodeoxynucleotide (ssODN) (Table S3) harbouring the Flag-HA sequence flanked by homology sequences to ZFC3H1 around the cleavage site. Single cells were isolated and amplified. HEK-293T clones expressing Flag-HA ZFC3H1 were identified by PCR and confirmed by sequencing as well as Western blot using anti-HA and anti-Flag antibodies.

### RNAi

Production of short hairpin RNA (shRNA)-expressing lentiviral particles was performed as described previously^53^ using plasmids expressing shRNAs targeting ZFC3H1 (Sigma-Aldrich MISSION shRNA, TRCN0000130498) or a non-targeting control (Addgene, plasmid 1864). For knockdown experiments, HeLa cells were either transduced with lentiviral particles and harvested 5 days later, or transfected with siRNAs shown in Table S4 using Interferin (PolyPlus) and harvested 72 h later, as described previously^54^.

### ZFC3H1 protein complex purification

ZFC3H1 complex was purified from Dignam high salt nuclear extracts (https://dx.doi.org/10.17504/protocols.io.kh2ct8e) from HEK-293T cells stably expressing Flag-HA-ZFC3H1 by two-step affinity chromatography (<u>10.17504/protocols.io.kgrctv6</u>). Sequential Flag and HA immunoprecipitations were performed on equal amounts of proteins. Silver staining was performed according to the manufacturer’s instructions (SilverQuest, Invitrogen). Elutions were precipitated using ProteoExtract Protein precipitation kit (Millipore) according to the manufacturer’s instructions.

Mass spectrometry was performed at the Taplin Facility, Harvard University, Boston, MA. Briefly, precipitates were resuspended in 50 μl ammonium bicarbonate solution (50 mM) with 10% acetonitrile by gentle vortexing. Ten μl of modified sequencing-grade trypsin (20 ng/ μl; Promega, Madison, WI) was added and samples were then placed in a 37°C room overnight. Samples were acidified with 5 μl of formic acid solution (20%) and then desalted by STAGE tip^55^.

On the day of analysis, the samples were reconstituted in 5 - 10 µl of HPLC solvent A (2.5% acetonitrile, 0.1% formic acid). A nano-scale reverse-phase HPLC capillary column was created by packing 2.6 µm C18 spherical silica beads into a fused silica capillary (100 µm inner diameter x ∼30 cm length) with a flame-drawn tip^56^. After equilibrating the column, each sample was loaded onto the column using a Famos auto sampler (LC Packings, San Francisco CA). A gradient was formed and peptides were eluted with increasing concentrations of solvent B (97.5% acetonitrile, 0.1% formic acid).

As peptides eluted, they were subjected to electrospray ionization and then entered into an LTQ Orbitrap Velos Pro ion-trap mass spectrometer (Thermo Fisher Scientific, Waltham, MA). Peptides were detected, isolated, and fragmented to produce a tandem mass spectrum of specific fragment ions for each peptide. Peptide sequences were determined by matching protein databases with the acquired fragmentation pattern by the software program, Sequest (Thermo Fisher Scientific, Waltham, MA)^57^. All databases include a reversed version of all the sequences and the data was filtered to between a one and two percent peptide false discovery rate.

### Coimmunoprecipitation analysis

Coimmunoprecipitation was performed using nuclear extracts of HeLa cells. Cells were lysed in ice-cold hypotonic buffer [20 mM tris (pH 7.6), 10 mM KCl, and 1.5 mM MgCl2] supplemented with EDTA-free complete protease inhibitor mixture (Roche) for 15 min on ice. NP-40 was added at 0.5% final, and extracts were centrifuged 1 min at 14,000g/4°C. The pellet (nuclei) was resuspended in nuclease buffer [20 mM tris (pH 7.6), 150 mM NaCl, 1.5 mM MgCl2, 2.5 mM CaCl2, and 0.5 μl of 100 mM phenylmethylsulfonyl fluoride] and incubated with micrococcal nuclease (2 × 103 U/ml; New England Biolabs) for 2 hours at 4°C. Lysates were cleared by centrifugation at 14,000g/4°C for 10 min and diluted in immunoprecipitation buffer [50 mM tris (pH 7.6), 150 mM NaCl, and 1% NP-40] supplemented with protease inhibitors. Protein concentration was determined using the Bradford reagent (Bio-Rad). Immunoprecipitations were performed using 400 μg of protein extracts with the indicated antibodies (2 μg) and rotated overnight at 4°C. Protein A Dynabeads were washed three times in immunoprecipitation buffer, added to protein extracts/antibody solution, and incubated for 2 hours at 4°C. Immunoprecipitates were washed extensively with the immunoprecipitation buffer. Where indicated, samples were incubated with RNAse A/T cocktail (Invitrogen AM2286; 1.2 μl/ml of immunoprecipitation buffer) for 30 min at RT on a rotating wheel, followed by 5 washes with the immunoprecipitation buffer. Samples were resuspended in protein sample loading buffer, boiled for 5 min, and analyzed by Western blotting using the antibodies shown in Table S2.

### Cell fractionation analysis

HeLa cells were seeded in 150 mm culture dishes the day prior to protein extraction. Cytoplasmic proteins were extracted using a mild lysis buffer (10 mM Hepes pH 7.9, 10 mM KCl, 0.1 mM EDTA pH 8.0, 2 mM MgCl2, 1 mM DTT, EDTA-free protease and phosphatase inhibitor). The cell pellet was incubated for 10 min on ice, adding 0.07% NP-40 and incubating for an additional 10 min on ice. After centrifugation (3000 rpm, 5min, 4°C), cytoplasmic fraction was collected. The pellet was washed with ice cold PBS supplemented with protease and phosphatase inhibitors. After centrifugation (3000 rpm, 5 min 4°C), the supernatant was discarded leaving the packed nuclear volume (PNV). For extraction of nuclear soluble proteins, nuclei were resuspended drop-wise in 1 x PNV of hypotonic buffer (20 mM Hepes pH 7.9, 20 mM NaCl, 1 mM EDTA pH 8.0, 1.5 mM MgCl2, 10% glycerol, 1 mM DTT, EDTA-free protease and phosphatase inhibitor), followed by the addition of 1 x PNV of high salt buffer (20 mM Hepes pH 7.9, 800 mM NaCl, 1 mM EDTA pH 8.0, 1.5 mM MgCl2, 10% glycerol, 1 mM DTT, EDTA-free protease and phosphatase inhibitor). Tubes with the samples were rotated on a wheel for 20 min at 4°C, followed by centrifugation (20 min 14 000 rpm, 4°C). Supernatant containing the nuclear soluble fraction was collected. The pellet was measured and chromatin bound proteins were extracted by adding 2 volumes of mild salt buffer (20 mM Hepes pH 7.9, 150 mM NaCl, 1 mM EDTA pH 8.0, 1.5 mM MgCl2, 10% glycerol, 1 mM DTT, EDTA-free protease and phosphatase inhibitor) together with 250 U Benzonase (Sigma Aldrich) per ml. Samples were incubated for 15 min at 37°C on a rotating wheel. After centrifugation (15 min, 14 000 rpm, 4°C), the chromatin-bound fraction was collected. Proteins were analyzed by Western blotting using the antibodies shown in Table S2.

### Quantitative RT-PCR

Total RNA was extracted from HeLa cells using TRIzol (ThermoFisher Scientific) according to the manufacturer’s instructions. Extracts were treated with DNase I (Promega) and reverse transcribed using SuperScript III First-Strand Synthesis System (ThermoFisher Scientific). RT products were amplified by real time PCR (LightCycler™ 480, Roche) using SYBR Green I Master mix (Roche) with the indicated oligonucleotides. Q-PCR cycling conditions are available on request. Sequences of qPCR primers used in this study are shown in Table S3.

### RNA Immunoprecipitation

RIP was performed as previously described^58^. Briefly, HeLa cells were seeded in 100 mm culture dishes and incubated overnight at 37°C. Cells were lysed for 10 min in RIP buffer (20 mM HEPES, pH 7.5, 150 mM NaCl, 2.5 mM MgCl_2_•6H_2_O, 250 mM sucrose, 0.05% (v/v) NP-40 and 0.5% (v/v) Triton X-100) containing 20 U ml^−1^ of RNasin (Promega), 1 mM DTT, 0.1 mM PMSF and EDTA-free protease and phosphatase inhibitor. After centrifugation, lysates were incubated for overnight at 4°C with 2 μg of antibodies recognizing MTR4 or IgG control and then incubated for 1 h at 4°C with Dynabeads™ Protein A (ThermoFisher Scientific). After incubation, beads were washed five times with RIP buffer for 5 min at 4°C and RNA was extracted using TRIzol (Thermo Fisher Scientific) according to the manufacturer’s instructions. RNA was treated with DNAse I (Promega) and RT was performed using SuperScript™ III Reverse Transcriptase (ThermoFisher Scientific) according to the manufacturer’s instructions. cDNAs were used to perform qPCRs using LightCycler™ 480 SYBR Green I Master mix (Roche), according to the manufacturer’s instructions, using the primers shown in Supplementary Table S3.

### RNA-seq

For RNA-seq, total RNA was extracted from HeLa cells using TRIzol (Thermo Fisher Scientific) according to the manufacturer’s instructions. RNA-seq (paired end, 125 bp) was carried out by Novogene in triplicates.

### ChIP, library preparation, and sequencing

Chromatin immunoprecipitation followed by high throughput sequencing (ChIP-seq)^59^ was performed from HeLa cells using the ChIP-IT High Sensitivity Kit from Active Motif (reference 53040) according to the manufacturer’s instructions, as described previously^53^. Sonication was performed using the Qsonica Q700 Sonicator with microtip of 1/8 inches (reference 4418) at 11% amplitude and 13 min of processing time (30-s “ON” and 30-s “OFF”). Each ChIP used 30 μg of chromatin together with 4 μg of antibody detecting ZFC3H1, RBM26, PAPγ or RNAPII (Supplementary Table S2). ChIP-seq libraries were constructed using the Next Gen DNA Library Kit (Active Motif, 53216 and 53264). Library quality was assessed using Agilent 2100 Bioanalyzer and Agilent High Sensitivity DNA assay. Libraries were sequenced with 2^nd^ generation sequencing chemistry on a Nextseq500 (Illumina) at the GENOM’IC facility, Institut Cochin, Paris.

### ChIP and Re-ChIP–qPCR

ZFC3H1 and PAPγ ChIP were performed using the iDeal ChIP-qPCR Kit (Diagenode, catalog no. C01010180) following the manufacturer’s instructions, as described previously^53^. HeLa cells were sonicated using the Bioruptor Pico (Diagenode, catalog no. B01060001) for 8 cycles of 30-s ON and 30-s OFF at high-power setting. ChIP was performed using the ChIP-IT Express Enzymatic Kit (Active Motif, catalog no. 53009), following the manufacturer’s instructions. Chromatin was digested for 7 min, and 50 μg of chromatin and 3 μg of antibody were used.

Re-ChIP was performed using the Re-ChIP-IT Kit (Active Motif, catalog no. 53016) according to the manufacturer’s instruction. Chromatin for re-ChIP was prepared using the ChIP-IT High Sensitivity Kit (Active Motif, catalog no. 53040), as described above. For each Re-ChIP, 50 μg of chromatin and 3 μg of antibody were used. Antibodies and sequences of primers used for real-time qPCR analysis are shown in Tables S2 and S3.

### PAT assay

PAT assays were performed as previously described^60,61^. Briefly, total RNA was extracted from cells using TRIzol (ThermoFisher Scientific) according to the manufacturer’s instructions. RNA was treated with RQ1 DNase (Promega) in the presence of RNasin (Promega), then reverse transcribed using SuperScript™ III (Invitrogen™) using either oligo (dT)-anchor (5’-GCGAGCTCCGCGGCCGCGTTTTTTTTTTTT-3’) or random hexamer primer. For normalization, cDNAs synthesized using random hexamer priming were amplified by quantitative PCR using the Forward and Reverse oligonucleotide pairs indicated in Table S3. Abundance of PCR products was calculated relative to the siCon sample, which was attributed a value of 1. Quantitative PCR reactions, using normalized amounts of cDNA generated using oligo (dT)-anchor priming, were then performed using the forward primer indicated in Table S3 together with oligo(dT)-anchor (50 μM). PCR products were run on 5% non-denaturing polyacrylamide TAE gels, stained with SYBR® Gold and visualized with ChemiDoc MP High-end imaging system.

Where indicated, samples were treated with RNase H. Briefly, DNase-treated RNA was incubated for 15 minutes at 70°C with 2 μL of Oligo(dT)20 in a thermocycler and the mixture was slowly cooled down to room temperature. Samples were then incubated for 1 hour at 37°C with 4 U RNase H (Invitrogen™) or mock-treated as a control prior to reverse transcription using oligo (dT) primer and qPCR amplification as described above.

### Bioinformatic analyses

For analysis of ChIP-seq data, sequencing reads were first filtered, using fastq_illumina_filter, and quality control of filtered reads was performed using FastQC. Filtered reads were then aligned onto the HG38 genome^62^ using the Burrows-Wheeler Aligner^63^ with default parameters. The sorted BAM files generated by SAMtools^64^ keeping only reads with a mapping quality at least 30 were then normalized by DeepTools^65^ bamCoverage function, with a bin size of 10 bp. RPGC normalization was applied, with an effective genome size of 2,913,022,398 bp according to DeepTools’ user manual instructions. Peak calling was performed using NormR’s enrichR function, searching for enrichment of each BAM file of ChIP-seq reads against the input BAM file using a false discovery rate threshold of 1e-4. Genomic Ranges^66^ was then used to determine overlap between the peak range and genomic features of interest, such as genes with a TSS and TES from GRCh38 and enhancers in HeLa from ENCODE. Profile matrices were extracted from the normalized data files using DeepTools’ computeMatrix using a bin size of 10 bp. Profiles were generated +/−5 kbp of TSSs and quantification of normalized reads was performed on +/−500 bp surrounding TSSs. Genomic elements and protein coding genes were obtained from Ensembl. Average profiles around genomic TSS were generated using SeqPlots^67^. Heatmaps were generated based on the indicated features with genomation (DOI: 10.18129/B9.bioc.genomation) using the gridHeat function, as performed on profile matrices generated by DeepTools. Proportional Venn diagrams were plotted using ‘Vennerable’ R package (https://github.com/js229/Vennerable).

For expression analyses, RNA-seq data were obtained from GSE131255^25^ or the present study. Forward and reverse RNA-seq reads were filtered using Ensembl reference coding genes to extract only antisense transcripts. Heatmaps were generated with genomation, as for ChIP-seq.

### Statistical analysis

Data presented as histograms are shown as means ± SEM. Comparison between two groups was analyzed by two-tailed Student’s t test, and asterisks represented significance defined as *P < 0.05, **P < 0.01, or ***P < 0.001. Enrichments in Venn diagrams were performed using Fisher’s exact test. Comparison of ChIP-seq signal in box plots was performed using the Wilcoxon pair-wise test.

### Data Availability

ChIP-seq and RNA-seq data have been deposited at GEO (GSE189157). Mass spectrometry data have been deposited at Massive (MSV000089482).

## Acknowledgements

The work carried out in this study was supported by ERC CoG RNAmedTGS, MSD Hide Inflame & Seq, SIDACTION and ANRS to RK, FRM (DEQ34940) and ANRS (#229035) to OC. CA was supported by a fellowship from ANRS, DD by scholarships from La Ligue Nationale Contre le Cancer (LNCC) and ANRS, KS by a fellowship from ARC (PDF20160604491). We thank Aymeric Chartier for help with PAT assays.

## Author contributions

XC prepared proteomic samples; XC, MN, CA and MS performed immunoblotting, fractionation analysis and RIP; CA and KS performed RT-qPCR; MH performed ChIP-seq; ML and MS performed ChIP- and re-ChIP-qPCR; CA and MS performed RNA-seq; CA performed PAT assay; DD performed data analysis in consultation with AH and OC; RK conceived the project; OC and RK supervised the project; XC, OC and RK wrote the paper with input from all authors.

## Additional Information

Correspondence and requests for materials should be addressed to Rosemary Kiernan.

**Supplementary Fig 1. Identification of the endogenous ZFC3H1 complex. A** Extracts of HEK-293T expressing Flag-HA-tagged ZFC3H1 (FH-ZFC3H1) were immunoprecipitated using anti-HA or anti-Flag antibodies, as indicated. Immunoprecipitates were blotted with anti-ZFC3H1 or anti-MTR4 antibodies, as indicated. **B** SDS-PAGE analysis followed by silver staining of eluates of tandem affinity purified nuclear extracts from 293T stably expressing Flag-HA-ZFC3H1 or control cells. **C** Representative gene ontology pathways (biological pathways) of ZFC3H1-associated proteins identified by mass spectrometry (n=131; 2 or more peptides, FC>2).

**Supplementary Fig 2 Interactions between different PAXT subunits are mostly RNA-independent. A** Co-immunoprecipitation analysis of ZFC3H1 interactome. HeLa nuclear extracts were immunoprecipitated using an IgG control or antibodies indicated on the figure above the blots. Immunoprecipitates and an aliquot of nuclear extract (input, 5%) were analyzed by SDS-PAGE followed by immunoblot using antibodies against ZFC3H1, RBM26, RBM27, YTHDC1, YTHDC2, PABPN1 or PAPγ, as indicated. **B** HeLa nuclear extracts were immunoprecipitated using antibodies against ZFC3H1, PAPγ or an IgG control and treated with RNase A/T or mock-treated, as indicated on the figure. Immunoprecipitates and an aliquot of nuclear extract (input, 5%) were analyzed by SDS-PAGE followed by immunoblot using the antibodies indicated on the right of the blots.

**Supplementary Fig 3 PAXT subunits are associated with chromatin. A** Cytosolic (cyto), nucleoplasmic (nuc) and chromatin (chrom) extracts of HeLa cells were analyzed by immunoblotting using the indicated antibodies. **B** Distribution of ChIP-seq peaks of RNAPII, ZFC3H1, RBM26 or PAPγ at transcription start site (TSS), gene body (GB), transcript end site (TES), enhancers (Enh) or other regions (Other). Number of peaks expressed as a percentage of the total peaks is shown at the left of the histograms, and total numbers of peaks are shown below. **C** Browser shots of RNAPII, ZFC3H1, RBM26, and PAPγ ChIP-seq signal over representative genes in HeLa cells. A schematic representation of the gene is shown above. **D** Anti-PAPγ specifically recognizes PAPγ. Occupancy of PAPγ at the TSS of the indicated genes detected by ChIP-qPCR using PAPγ antibodies (24284-1 and A302-426A, respectively) and chromatin from Hela cells transfected with siRNAs targeting PAPγ or a non-targeting control, as indicated. Data represent mean ± SEM obtained from 3 independent experiments (\*\*\**P* < 0.001\*\**P* < 0.01, \**P* < 0.05, independent Student’s *t* test).

**Supplementary Fig 4 PAXT subunits are co-recruited to chromatin. A** ChIP-qPCR showing localization of ZFC3H1 and PAPγ at the TSS region of the indicated genes. Data represent mean ± SEM obtained from 3 independent experiments (\*\*\**P* < 0.001, independent Student’s *t* test). **B** Scatter plots showing the normalized ChIP-seq signal of ZFC3H1, RBM26 and PAPγ relative to that of RNAPII at the TSS of genes with a high or low occupancy of ZFC3H1 at the TSS, as indicated.

**Supplementary Fig 5 Ablation of ZFC3H1 by specific shRNA.** HeLa cells transduced with lentiviral particles expressing shRNA against ZFC3H1 or a non-targeting control were harvested 7 days post-infection. Extracts were analyzed by immunoblot using the indicated antibodies.

**Supplementary Fig 6. Loss of PAP**g **stabilizes PROMPTs genome-wide. A** HeLa cells transfected with siRNA against PAPγ or a non-targeting control were harvested 4 days post-infection. Extracts were analyzed by immunoblot using the indicated antibodies. **B** Volcano plot showing the differential expression of mRNAs in extracts of HeLa cells transfected with siCon or siPAPγ.

**Supplementary Fig 7 Loss of PAXT subunits stabilizes PROMPTs. A** Immunoblot analysis of HeLa cell extracts following transfection with siRNAs directed against MTR4, ZFC3H1, PAPγ, or a control (con), as indicated on the figure. **B** Immunoblot analysis of HeLa cell extracts following transfection with siMTR4#2, siZFC3H1#2, siPAPγ#2, or a control (con#2), as indicated on the figure. **C** Total RNA extracts of HeLa cells transfected with the siRNAs shown in B were analyzed by RT-q-PCR using the oligonucleotide pairs indicated and the values were normalized to those for the control transfection, which was attributed a value of 1. Data represent mean ± SEM obtained from 3 independent experiments (\*\*\**P* < 0.001, \*\**P* < 0.01, \**P* < 0.05, independent Student’s *t* test). **D, E** An aliquot of HeLa cell extract (input) and immunoprecipitates (IP) obtained using control IgG, anti-MTR4 (D) or anti-PAPγ (E) antibodies, as indicated, and flow through samples (FT) were analyzed by Western blot using the indicated antibodies.

## References

1. Core, L.J., Waterfall, J.J. & Lis, J.T. Nascent RNA sequencing reveals widespread pausing and divergent initiation at human promoters. Science 322, 1845–1848 (2008).

2. Neil, H. et al. Widespread bidirectional promoters are the major source of cryptic transcripts in yeast. Nature 457, 1038–1042 (2009).

3. Preker, P. et al. RNA exosome depletion reveals transcription upstream of active human promoters. Science 322, 1851–1854 (2008).

4. Ogami, K. et al. An Mtr4/ZFC3H1 complex facilitates turnover of unstable nuclear RNAs to prevent their cytoplasmic transport and global translational repression. Genes Dev. 31, 1257–1271 (2017).

5. Yang, F. et al. Shape of promoter antisense RNAs regulates ligand-induced transcription activation. Nature 595, 444–449 (2021).

6. Fasken, M.B. et al. Insight into the RNA Exosome Complex Through Modeling Pontocerebellar Hypoplasia Type 1b Disease Mutations in Yeast. Genetics 205, 221–237 (2017).

7. Mitchell, P., Petfalski, E., Shevchenko, A., Mann, M. & Tollervey, D. The exosome: a conserved eukaryotic RNA processing complex containing multiple 3’-->5’ exoribonucleases. Cell 91, 457–466 (1997).

8. Schmid, M. & Jensen, T.H. Controlling nuclear RNA levels. Nat. Rev. Genet. 19, 518–529 (2018).

9. Zinder, J.C. & Lima, C.D. Targeting RNA for processing or destruction by the eukaryotic RNA exosome and its cofactors. Genes Dev. 31, 88–100 (2017).

10. Hernandez, H., Dziembowski, A., Taverner, T., Seraphin, B. & Robinson, C.V. Subunit architecture of multimeric complexes isolated directly from cells. EMBO Rep. 7, 605–610 (2006).

11. Liu, Q., Greimann, J.C. & Lima, C.D. Reconstitution, activities, and structure of the eukaryotic RNA exosome. Cell 127, 1223–1237 (2006).

12. Allmang, C. et al. The yeast exosome and human PM-Scl are related complexes of 3’ --> 5’ exonucleases. Genes Dev. 13, 2148–2158 (1999).

13. Januszyk, K., Liu, Q. & Lima, C.D. Activities of human RRP6 and structure of the human RRP6 catalytic domain. RNA 17, 1566–1577 (2011).

14. Staals, R.H. et al. Dis3-like 1: a novel exoribonuclease associated with the human exosome. EMBO J. 29, 2358–2367 (2010).

15. Stead, J.A., Costello, J.L., Livingstone, M.J. & Mitchell, P. The PMC2NT domain of the catalytic exosome subunit Rrp6p provides the interface for binding with its cofactor Rrp47p, a nucleic acid-binding protein. Nucleic Acids Res 35, 5556–5567 (2007).

16. Tomecki, R. et al. The human core exosome interacts with differentially localized processive RNases: hDIS3 and hDIS3L. EMBO J. 29, 2342–2357 (2010).

17. Zinder, J.C., Wasmuth, E.V. & Lima, C.D. Nuclear RNA Exosome at 3.1 A Reveals Substrate Specificities, RNA Paths, and Allosteric Inhibition of Rrp44/Dis3. Mol. Cell 64, 734–745 (2016).

18. Bernstein, J., Patterson, D.N., Wilson, G.M. & Toth, E.A. Characterization of the essential activities of Saccharomyces cerevisiae Mtr4p, a 3’->5’ helicase partner of the nuclear exosome. J. Biol. Chem. 283, 4930–4942 (2008).

19. LaCava, J. et al. RNA degradation by the exosome is promoted by a nuclear polyadenylation complex. Cell 121, 713–724 (2005).

20. Schilders, G., van Dijk, E. & Pruijn, G.J. C1D and hMtr4p associate with the human exosome subunit PM/Scl-100 and are involved in pre-rRNA processing. Nucleic Acids Res 35, 2564–2572 (2007).

21. Lubas, M. et al. The human nuclear exosome targeting complex is loaded onto newly synthesized RNA to direct early ribonucleolysis. Cell Rep. 10, 178–192 (2015).

22. Lubas, M. et al. Interaction profiling identifies the human nuclear exosome targeting complex. Mol. Cell 43, 624–637 (2011).

23. Meola, N. et al. Identification of a Nuclear Exosome Decay Pathway for Processed Transcripts. Mol. Cell 64, 520–533 (2016).

24. Silla, T., Karadoulama, E., Makosa, D., Lubas, M. & Jensen, T.H. The RNA Exosome Adaptor ZFC3H1 Functionally Competes with Nuclear Export Activity to Retain Target Transcripts. Cell Rep. 23, 2199–2210 (2018).

25. Silla, T. et al. The human ZC3H3 and RBM26/27 proteins are critical for PAXT-mediated nuclear RNA decay. Nucleic Acids Res. 48, 2518–2530 (2020).

26. Bresson, S.M. & Conrad, N.K. The human nuclear poly(a)-binding protein promotes RNA hyperadenylation and decay. PLoS Genet 9, e1003893 (2013).

27. Bresson, S.M., Hunter, O.V., Hunter, A.C. & Conrad, N.K. Canonical Poly(A) Polymerase Activity Promotes the Decay of a Wide Variety of Mammalian Nuclear RNAs. PLoS Genet 11, e1005610 (2015).

28. Juge, F., Zaessinger, S., Temme, C., Wahle, E. & Simonelig, M. Control of poly(A) polymerase level is essential to cytoplasmic polyadenylation and early development in Drosophila. EMBO J. 21, 6603–6613 (2002).

29. Kuhn, U., Buschmann, J. & Wahle, E. The nuclear poly(A) binding protein of mammals, but not of fission yeast, participates in mRNA polyadenylation. RNA 23, 473–482 (2017).

30. Lemay, J.F. et al. The Nuclear Poly(A)-Binding Protein Interacts with the Exosome to Promote Synthesis of Noncoding Small Nucleolar RNAs. Mol. Cell 37, 34–45.

31. Kyriakopoulou, C.B., Nordvarg, H. & Virtanen, A. A novel nuclear human poly(A) polymerase (PAP), PAP gamma. J. Biol. Chem. 276, 33504–33511 (2001).

32. Perumal, K., Sinha, K., Henning, D. & Reddy, R. Purification, characterization, and cloning of the cDNA of human signal recognition particle RNA 3’-adenylating enzyme. J Biol Chem 276, 21791–21796 (2001).

33. Topalian, S.L. et al. Identification and functional characterization of neo-poly(A) polymerase, an RNA processing enzyme overexpressed in human tumors. Mol. Cell. Biol. 21, 5614–5623 (2001).

34. Yang, Q., Nausch, L.W., Martin, G., Keller, W. & Doublie, S. Crystal structure of human poly(A) polymerase gamma reveals a conserved catalytic core for canonical poly(A) polymerases. J. Mol. Biol. 426, 43–50 (2014).

35. Vanacova, S. et al. A new yeast poly(A) polymerase complex involved in RNA quality control. PLoS Biol. 3, e189 (2005).

36. Dobrev, N. et al. The zinc-finger protein Red1 orchestrates MTREC submodules and binds the Mtl1 helicase arch domain. Nat. Commun. 12, 3456 (2021).

37. Zhou, Y. et al. The fission yeast MTREC complex targets CUTs and unspliced pre-mRNAs to the nuclear exosome. Nat. Commun. 6, 7050 (2015).

38. Bentley, D.L. Coupling mRNA processing with transcription in time and space. Nat. Rev. Genet. 15, 163–175 (2014).

39. Bieberstein, N.I., Straube, K. & Neugebauer, K.M. Chromatin immunoprecipitation approaches to determine co-transcriptional nature of splicing. Methods Mol. Biol. 1126, 315–323 (2014).

40. Andersson, R. et al. An atlas of active enhancers across human cell types and tissues. Nature 507, 455–461 (2014).

41. Andersson, R. et al. Nuclear stability and transcriptional directionality separate functionally distinct RNA species. Nat. Commun. 5, 5336 (2014).

42. Passmore, L.A. & Coller, J. Roles of mRNA poly(A) tails in regulation of eukaryotic gene expression. Nat. Rev. Mol. Cell. Biol. 23, 93–106 (2022).

43. Belasco, J.G. All things must pass: contrasts and commonalities in eukaryotic and bacterial mRNA decay. Nat. Rev. Mol. Cell. Biol. 11, 467–478 (2010).

44. Hajnsdorf, E., Braun, F., Haugel-Nielsen, J. & Regnier, P. Polyadenylylation destabilizes the rpsO mRNA of Escherichia coli. Proc Natl Acad Sci U S A 92, 3973–3977 (1995).

45. Hajnsdorf, E. & Regnier, P. E. coli RpsO mRNA decay: RNase E processing at the beginning of the coding sequence stimulates poly(A)-dependent degradation of the mRNA. J Mol Biol 286, 1033–43 (1999).

46. Sarkar, N. Polyadenylation of mRNA in prokaryotes. Annu. Rev. Biochem. 66, 173–197 (1997).

47. Carpousis, A.J. The RNA degradosome of Escherichia coli: an mRNA-degrading machine assembled on RNase E. Annu Rev. Microbiol. 61, 71–87 (2007).

48. Mohanty, B.K. & Kushner, S.R. Bacterial/archaeal/organellar polyadenylation. Wiley Interdiscip Rev RNA 2, 256–276 (2011).

49. Houseley, J. & Tollervey, D. The many pathways of RNA degradation. Cell 136, 763–776 (2009).

50. Meola, N. & Jensen, T.H. Targeting the nuclear RNA exosome: Poly(A) binding proteins enter the stage. RNA Biol. 14, 820–826 (2017).

51. Soni, K. et al. Mechanistic insights into RNA surveillance by the canonical poly(A) polymerase Pla1 of the MTREC complex. Nat. Commun. 14, 772 (2023).

52. Almada, A.E., Wu, X., Kriz, A.J., Burge, C.B. & Sharp, P.A. Promoter directionality is controlled by U1 snRNP and polyadenylation signals. Nature 499, 360–363 (2013).

53. Salifou, K. et al. Chromatin-associated MRN complex protects highly transcribing genes from genomic instability. Sci Adv. 7(2021).

54. Contreras, X. et al. Nuclear RNA surveillance complexes silence HIV-1 transcription. PLoS Pathog 14, e1006950 (2018).

55. Rappsilber, J., Ishihama, Y. & Mann, M. Stop and go extraction tips for matrix-assisted laser desorption/ionization, nanoelectrospray, and LC/MS sample pretreatment in proteomics. Anal. Chem. 75, 663–670 (2003).

56. Peng, J. & Gygi, S.P. Proteomics: the move to mixtures. J. Mass Spectrom. 36, 1083–1091 (2001).

57. Eng, J.K., McCormack, A.L. & Yates, J.R. An approach to correlate tandem mass spectral data of peptides with amino acid sequences in a protein database. J. Am. Soc. Mass Spectrom. 5, 976–989 (1994).

58. Grasso, G. et al. NF90 modulates processing of a subset of human pri-miRNAs. Nucleic Acids Res. 48, 6874–6888 (2020).

59. Barski, A. et al. High-resolution profiling of histone methylations in the human genome. Cell 129, 823–837 (2007).

60. Chartier, A., Joly, W. & Simonelig, M. Measurement of mRNA Poly(A) Tail Lengths in Drosophila Female Germ Cells and Germ-Line Stem Cells. Methods Mol. Biol. 1463, 93–102 (2017).

61. Dufourt, J. et al. piRNAs and Aubergine cooperate with Wispy poly(A) polymerase to stabilize mRNAs in the germ plasm. Nat. Commun. 8, 1305 (2017).

62. Lander, E.S. et al. Initial sequencing and analysis of the human genome. Nature 409, 860–921 (2001).

63. Li, H. & Durbin, R. Fast and accurate short read alignment with Burrows-Wheeler transform. Bioinformatics 25, 1754–1760 (2009).

64. Bonfield, J.K. et al. HTSlib: C library for reading/writing high-throughput sequencing data. Gigascience 10(2021).

65. Ramirez, F. et al. deepTools2: a next generation web server for deep-sequencing data analysis. Nucleic Acids Res 44, W160–5 (2016).

66. Lawrence, M. et al. Software for computing and annotating genomic ranges. PLoS Comput Biol 9, e1003118 (2013).

67. Stempor, P. & Ahringer, J. SeqPlots - Interactive software for exploratory data analyses, pattern discovery and visualization in genomics. Wellcome Open Res 1, 14 (2016).

