## Supplementary Figures and Tables for "PAPγ associates with PAXT nuclear exosome to control the abundance of PROMPT ncRNAs": MTREC_supp figures_revision round 4v2.pdf

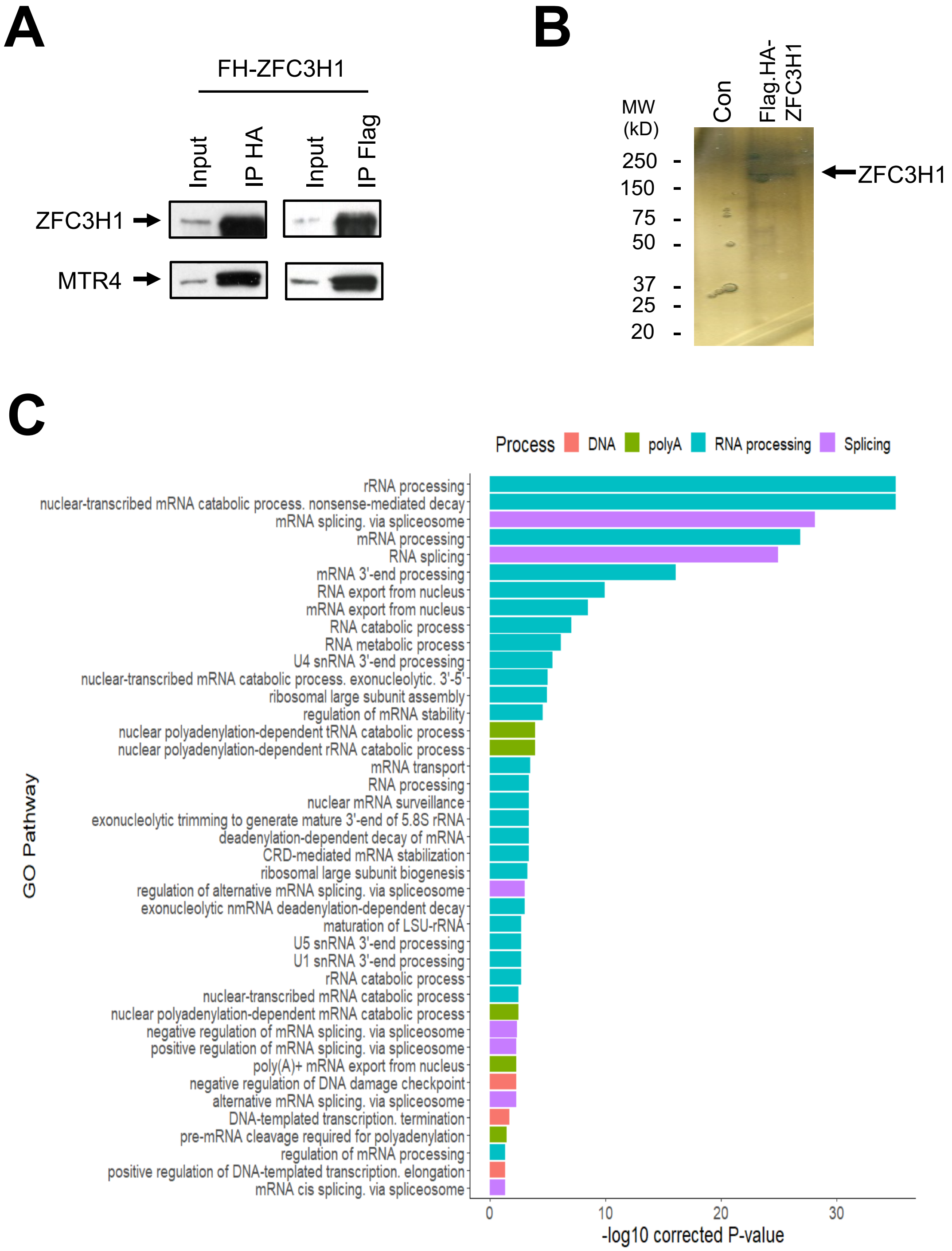


Supplementary Fig. 1

**Supplementary Fig 1. Characterization of the endogenous ZFC3H1 complex.** **A** Extracts of HEK-293T expressing Flag-HA-tagged ZFC3H1 (FH-ZFC3H1) were immunoprecipitated using anti-HA or anti-Flag antibodies, as indicated. Immunoprecipitates were blotted with anti-ZFC3H1 or anti-MTR4 antibodies, as indicated. **B** SDS-PAGE analysis followed by silver staining of eluates of tandem affinity purified nuclear extracts from 293T stably expressing Flag-HA-ZFC3H1 or control cells. **C** Representative gene ontology pathways (biological pathways) of ZFC3H1-associated proteins identified by mass spectrometry (n=131; 2 or more peptides, FC>2).

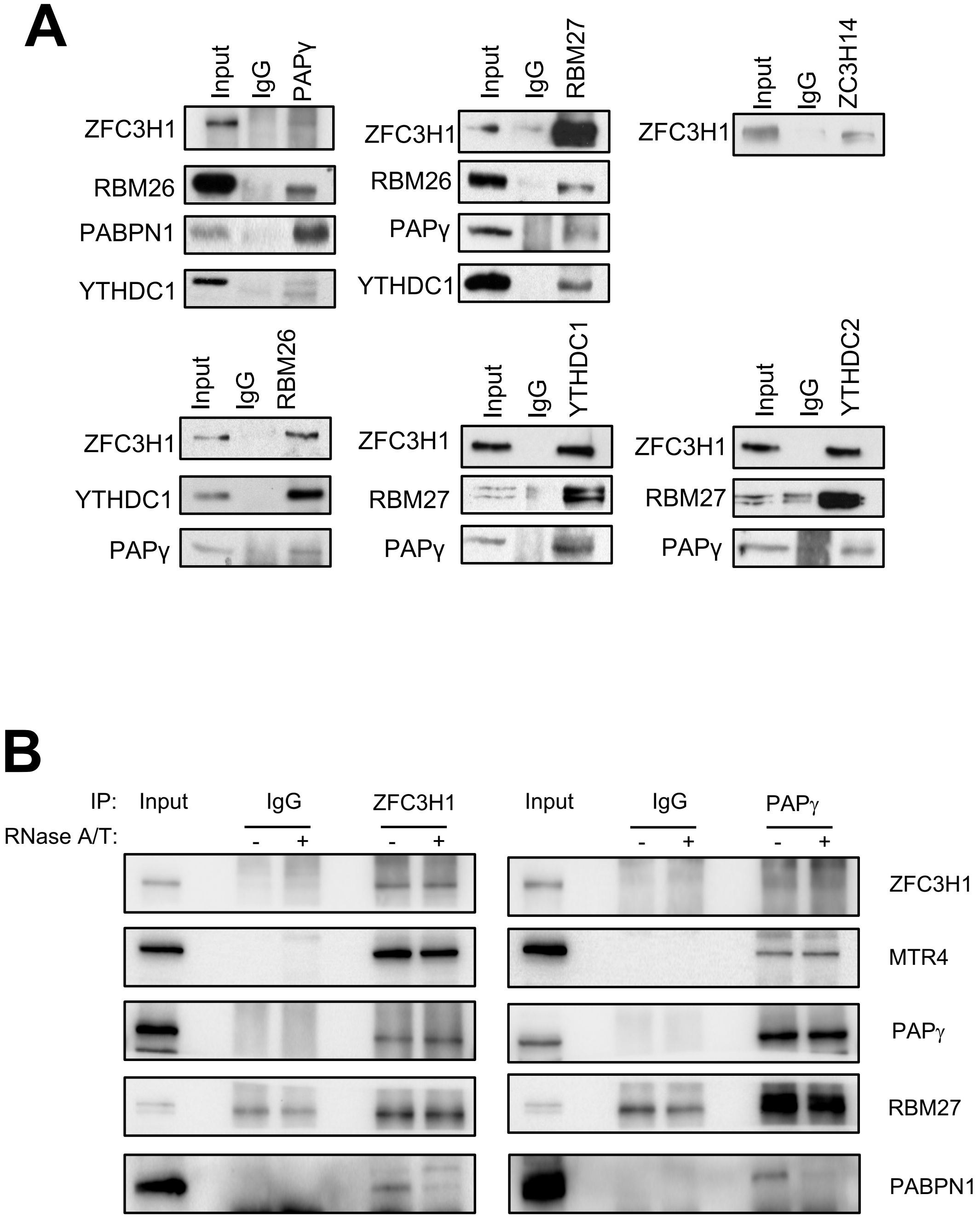


Supplementary Fig. 2

**Supplementary Fig 2 Interactions between different PAXT subunits are mostly RNA-independent. A** Co-immunoprecipitation analysis of ZFC3H1 interactome. HeLa nuclear extracts were immunoprecipitated using an IgG control or antibodies indicated on the figure above the blots. Immunoprecipitates and an aliquot of nuclear extract (input, 5%) were analyzed by SDS-PAGE followed by immunoblot using antibodies against ZFC3H1, RBM26, RBM27, YTHDC1, YTHDC2, PABPN1 or PAP$\gamma$, as indicated. **B** HeLa nuclear extracts were immunoprecipitated using antibodies against ZFC3H1, PAP$\gamma$ or an IgG control and treated with RNase A/T or mock-treated, as indicated on the figure. Immunoprecipitates and an aliquot of nuclear extract (input, 5%) were analyzed by SDS-PAGE followed by immunoblot using the antibodies indicated on the right of the blots.

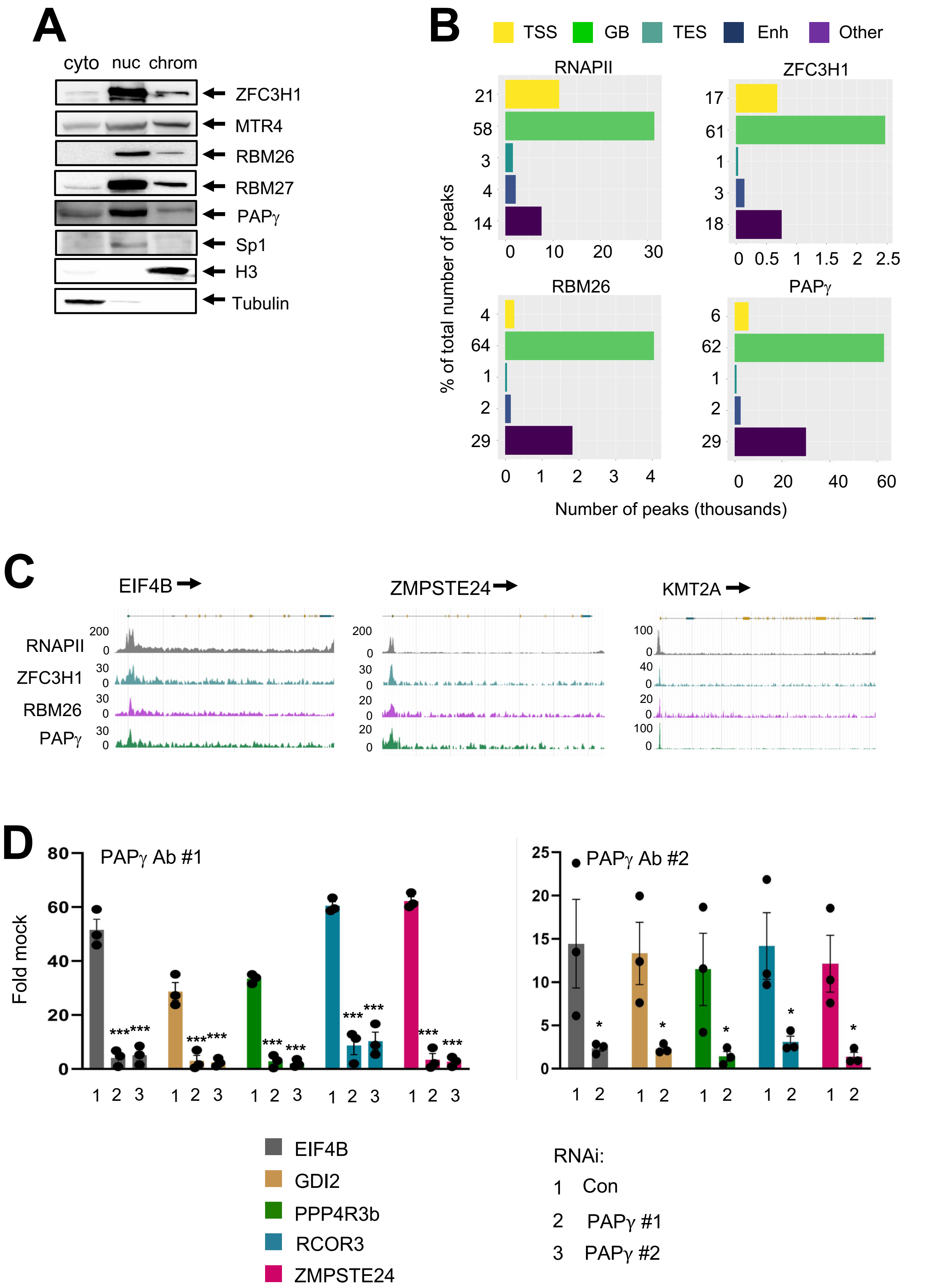


Supplementary Fig. 3

**Supplementary Fig 3 PAXT subunits are associated with chromatin. A** Cytosolic (cyto), nucleoplasmic (nuc) and chromatin (chrom) extracts of HeLa cells were analyzed by immunoblotting using the indicated antibodies. **B** Distribution of ChIP-seq peaks of RNAPII, ZFC3H1, RBM26 or PAPγ at transcription start site (TSS), gene body (GB), transcript end site (TES), enhancers (Enh) or other regions (Other). Number of peaks expressed as a percentage of the total peaks is shown at the left of the histograms, and total numbers of peaks are shown below. **C** Browser shots of RNAPII, ZFC3H1, RBM26, and PAPγ ChIP-seq signal over representative genes in HeLa cells. A schematic representation of the gene is shown above. **D** Anti-PAPγ specifically recognizes PAPγ. Occupancy of PAPγ at the TSS of the indicated genes detected by ChIP-qPCR using PAPγ antibodies (24284-1 and A302-426A, respectively) and chromatin from Hela cells transfected with siRNAs targeting PAPγ or a non-targeting control, as indicated. Data represent mean ± SEM obtained from 3 independent experiments (***$P < 0.001$**$P < 0.01$, *$P < 0.05$, independent Student's *t* test).

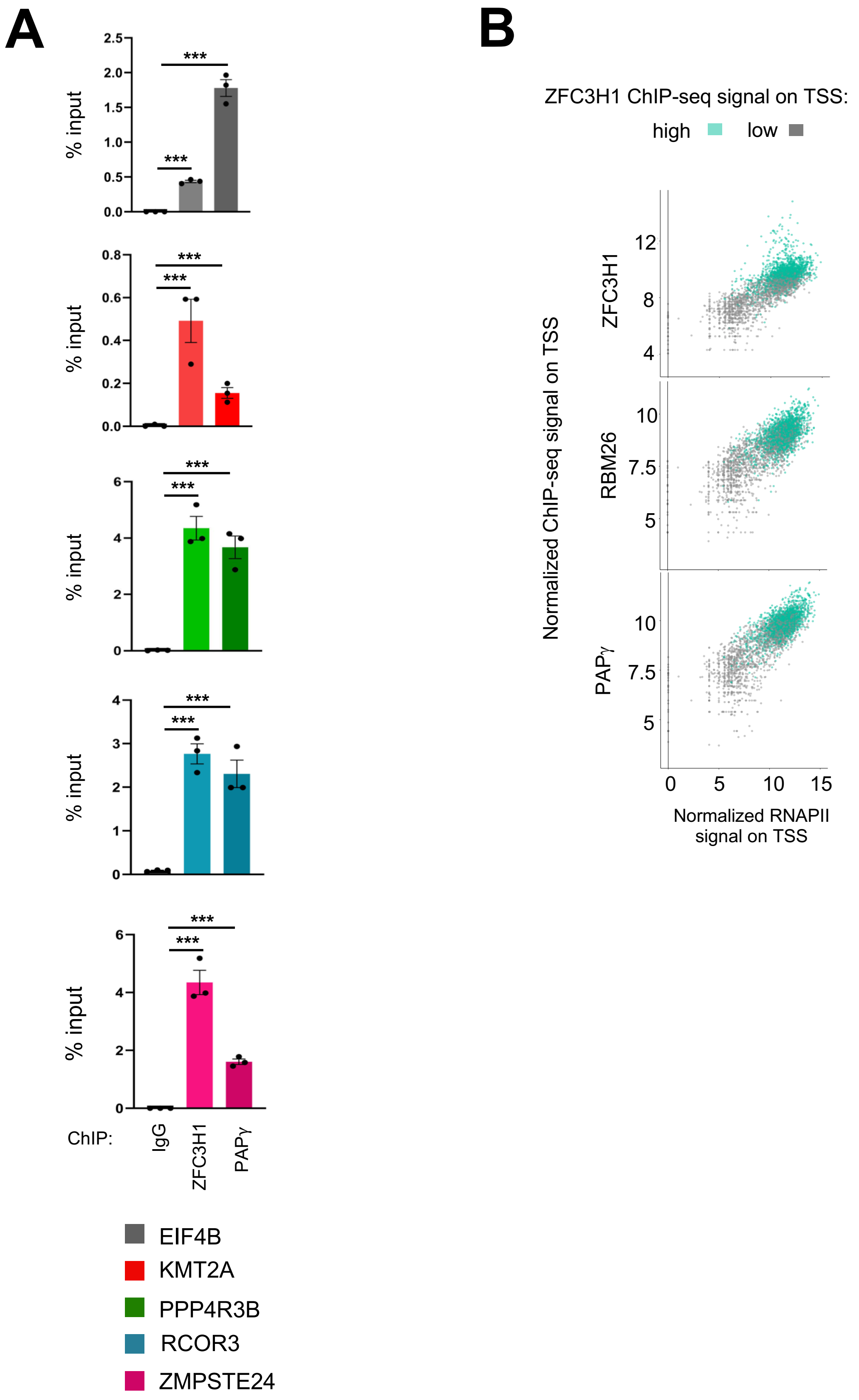


Supplementary Fig. 4

**Supplementary Fig 4 PAXT subunits are co-recruited to chromatin. A** ChIP-qPCR showing localization of ZFC3H1 and PAPγ at the TSS region of the indicated genes. Data represent mean ± SEM obtained from 3 independent experiments (****P* < 0.001, independent Student's *t* test). **B** Scatter plots showing the normalized ChIP-seq signal of ZFC3H1, RBM26 and PAPγ relative to that of RNAPII at the TSS of genes with a high or low occupancy of ZFC3H1 at the TSS, as indicated.

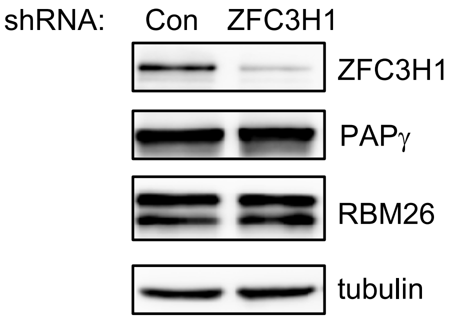


**Supplementary Fig 5 Ablation of ZFC3H1 by specific shRNA.** HeLa cells transduced with lentiviral particles expressing shRNA against ZFC3H1 or a non-targeting control were harvested 7 days post-infection. Extracts were analyzed by immunoblot using the indicated antibodies.

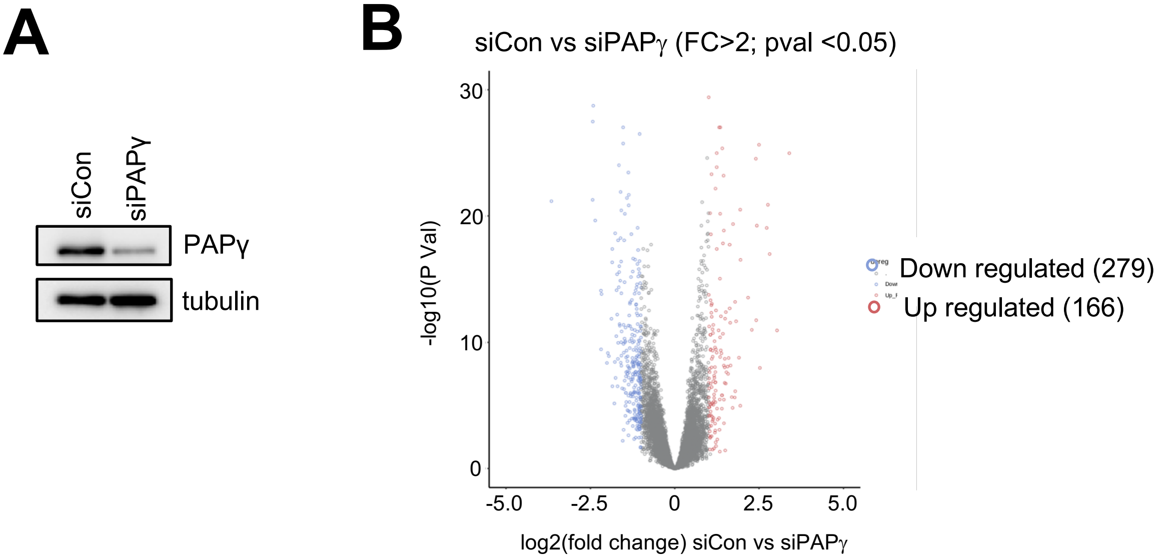


**Supplementary Fig 6. Loss of PAPγ modestly affects the expression of mRNAs. A** HeLa cells transfected with siRNA against PAPγ or a non-targeting control were harvested 4 days post-infection. Extracts were analyzed by immunoblot using the indicated antibodies. **B** Volcano plot showing the differential expression of mRNAs in extracts of HeLa cells transfected with siCon or siPAPγ.

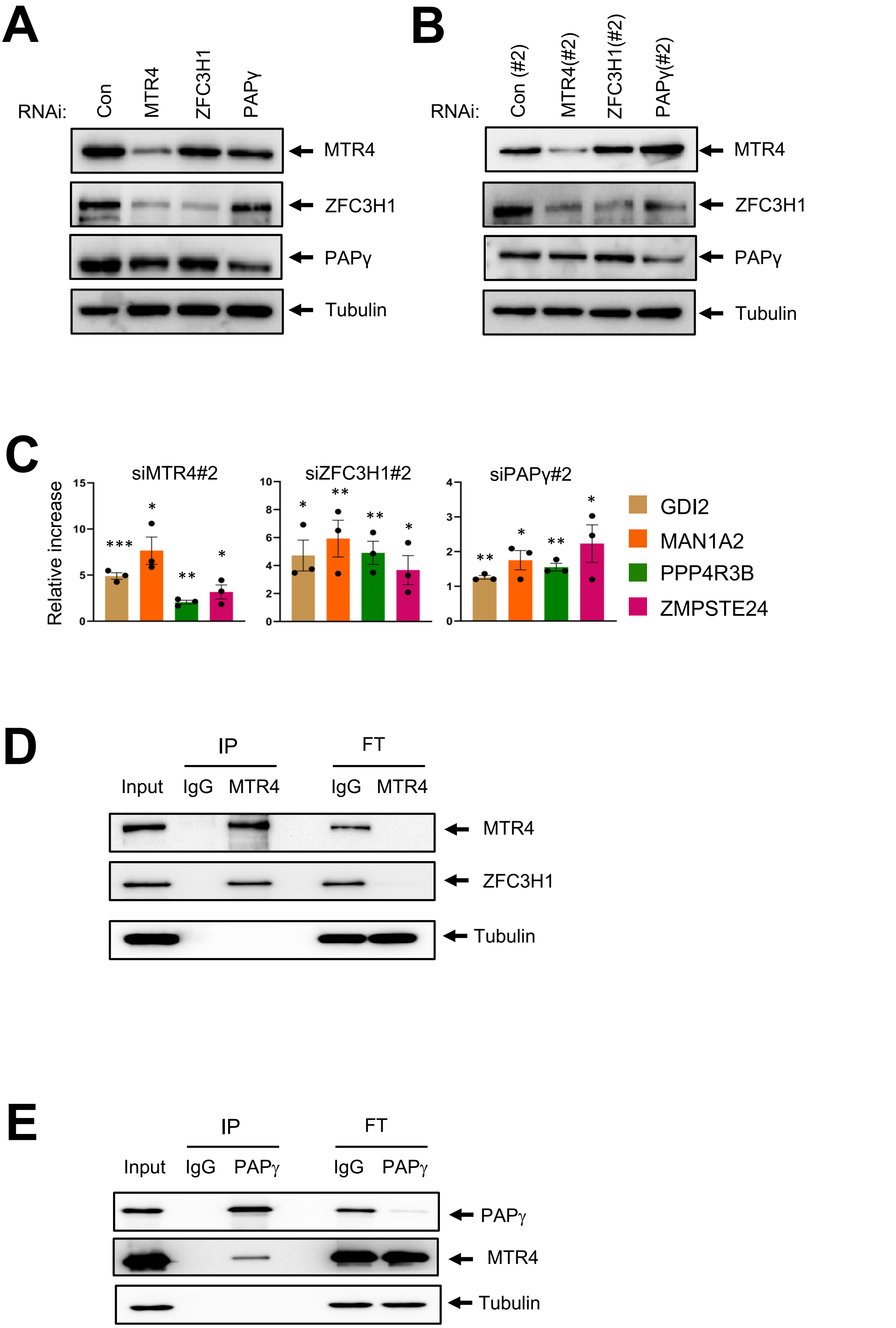


Supplementary Fig. 7

**Supplementary Fig 7 Loss of PAXT subunits stabilizes PROMPTs. A** Immunoblot analysis of HeLa cell extracts following transfection with siRNAs directed against MTR4, ZFC3H1, PAPγ, or a control (con), as indicated on the figure. **B** Immunoblot analysis of HeLa cell extracts following transfection with siMTR4#2, siZFC3H1#2, siPAPγ#2, or a control (con#2), as indicated on the figure. **C** Total RNA extracts of HeLa cells transfected with the siRNAs shown in B were analyzed by RT-q-PCR using the oligonucleotide pairs indicated and the values were normalized to those for the control transfection, which was attributed a value of 1. Data represent mean ± SEM obtained from 3 independent experiments (***$P$ < 0.001, **$P$ < 0.01, *$P$ < 0.05, independent Student's *t* test). **D, E** An aliquot of HeLa cell extract (input) and immunoprecipitates (IP) obtained using control IgG, anti-MTR4 (D) or anti-PAPγ (E) antibodies, as indicated, and flow-through samples (FT) were analyzed by Western blot using the indicated antibodies.

Table S2. Antibodies used in this study.

| Target | Reference | Supplier |
|---|---|---|
| CPSF6 | A301-356A | Bethyl Laboratories |
| Flag | M2, F1804 | Sigma-Aldrich |
| HA | SC-7392 | SCBT |
| Histone H3 | Ab1791 | Abcam |
| IgG mouse | SC-2025 | SCBT |
| IgG rabbit | SC-2027 | SCBT |
| MTR4 (IP) | ab70552 | Abcam |
| MTR4 (WB) | A300-614A | Bethyl Laboratories |
| MTR4 (RIP and WB) | 12719-2-AP | Proteintech |
| PABPN1 | 66807-1-Ig | Proteintech |
| PAPγ (WB ) | A302-427A | Bethyl Laboratories |
| PAPγ (IP, ChIP, ChIP-seq) | 24284-1-AP | Proteintech |
| PAPγ (IP, ChIP) | A302-426A | Bethyl Laboratories |
| RBM26 (IP, WB) | A301-215A | Bethyl Laboratories |
| RBM27 | A301-234A | Bethyl Laboratories |
| RNAPII | F12; SC-55492 | SCBT |
| Sp1 | SC-59 | SCBT |
| Tubulin | DM1A, T6199 | Sigma-Aldrich |
| YTHDC1 | A305-096A | Bethyl Laboratories |
| YTHDC2 | A303-025A | Bethyl Laboratories |
| ZC3H3 | A301-878A | Bethyl Laboratories |
| ZC3H14 | A303-992A | Bethyl Laboratories |
| ZFC3H1 | A301-457A | Bethyl Laboratories |

Table S3. Oligodeoxynucleotide primers used in this study.

| ssODN for genome editing | |
|---|---|
| Name | Sequence |
| ZFC3H1 -N ter | AATTGTCGGGCGGATCCCCGGACGGAGGGCTAAGGTTGTGTGGAAGGCGC TGCTCGCGAGATGGACTACAAAGACGATGACGACAAGCTCCATGGAGGATA CCCATACGATGTTCCAGATTACGCTGGAGGACTCGCGACCGCAGATACTCC GGCCCCGGCCTCCAGTGGCCTCTCGCCGAAGGAAGAAGGGGAG |

| Primers for RT-qPCR (PROMPTs) | | |
|---|---|---|
| Name | Forward | Reverse |
| EIF4B | GGCGAAATGGATGTGCTGAA | CGCCTGTAATCCCAGCTACT |
| GDI2 | GGTTTCATTGGGAAGCCGATC | GCTTCTGCTTACACCTGTGTC |
| KMT2A | AAACCTACGTCCCTCCACTG | GCGCTGTAACAACGGAATCT |
| MAN1A2 | GAGCGCGAAAGAAGAGAAGG | AGCCATTAACTCCGGTGACA |
| MRPL20 | TCTAGCGTGGGTGACAGATC | CATGCACTTGTGGTTCAGCT |
| PPP4R33 | CCCATCCCCGCTATATTCGT | ACACTGACCCACGTTGTAGT |
| RCOR3 | GGAGTGAGTGCTGTGGAGTG | ATAAACAGCCCCCAATGCCA |
| RPL8 | GCCAAGTGGTTGAGTGACCT | CCTGCAAACCATGACGTTCC |
| SLC4A1 | ACCTCTCCAACACAAGCTCA | CACAGCACTGCGATCTCTTC |
| ZMPSTE24 | AGGAGGCTAGGAAGGACTGA | CGTCCACTCTCTCATACGCT |

| Primers for ChIP and Re-ChIP | | |
|---|---|---|
| Name | Forward | Reverse |
| EIF4B | CTCTTTCCCTCTCCCAACAT | CGACTTCAGGAAATCAGTGG |
| GDI2 | TCTGTGCCCTCCTTGTTTAG | AAACCTGGCTTTCACTTTCA |
| KMT2A | CGAGGCCGCTATACAGATTG | GTGAAGTGAAGCAGCGAGAG |
| PPP4R3b | GTTGGATGGCACCAGTATGT | CTTCCTTTCTGTCCCTGGTT |
| RCOR3 | CAGAGAGGGCAAAGAGTCC | CCCAACCAAGCGCTAATAAA |
| ZMPSTE24 | GTTGATGCAACACAGCTCAG | CAGACACACTCTTCCCCTTG |

| Primers for PAT assay | | |
|---|---|---|
| Name | Forward | Reverse |
| PPP4R3B | AAAAGAACGTGGAGCCCTTT | GCCTGGAGAGTTGTGTCCAT |
| GDI2 | GGTTTCATTGGGAAGCCGATC | GCTTCTGCTTACACCTGTGTC |
| ZMPSTE24 | AGGAGGCTAGGAAGGACTGA | CGTCCACTCTCTCATACGCT |
| K-Ras | TGTATAGTGTAAACTGAAACA TGCAC | GTCACTGTAACTATTTTTATTAC |

Table S4. Sequence targeted by siRNAs used in this study.

| siRNAs | |
|---|---|
| **Target** | Sequence |
| **Control** | GCGCGCUUUGUAGGAUUCG(dTdT) |
| **Control #2** | UCUGCAAGGUUAGGCGUCU(dTdT) |
| **MTR4** | ACACUGAGCUGGAAAAUAA(dTdT) |
| **MTR4 #2** | AAGAAGUAUUCAGUAAUGC (dtdT) |
| **ZFC3H1** | CUAUAGCAGUCCAUCUUAA(dTdT) |
| **ZFC3H1 #2** | GUACAAUCGAUUCAUGAAA (dTdT) |
| **PAPγ** | ACAUCAAGAUGGCAUUAGA(dTdT) |
| **PAPγ #2** | UUUAUCUCAGGUUUCAAAG(dTdT) |
